# A prototype timsOmni platform enables confident annotation of key hypervariable regions of IgG immunoglobulins using low- and high-energy electron-based fragmentation

**DOI:** 10.1101/2025.10.29.685408

**Authors:** Simon Ollivier, Dina Schuster, Danique M.H. van Rijswijck, Iuliia Stroganova, Athanasios Smyrnakis, Jan Fiala, Mariangela Kosmopoulou, Detlev Suckau, Stuart Pengelley, Oliver Raether, Jean-François Greisch, Dimitris Papanastasiou, Albert J.R. Heck

## Abstract

The configuration of the first prototype timsOmni instrument, which integrates an Omnitrap linear ion trap into a timsTOF platform, is presented and applied to antibody analysis. A modified electrode design for the electron-based fragmentation (ExD) section of the Omnitrap platform is introduced, enhancing both the robustness and performance of the method. Optimal characterization of antibodies requires characterizing light and heavy chains as pairs in addition to sequencing their variable domains and identifying any modifications. This is best addressed using protein centric proteomics as heterogeneity information such as the specific clonal origin of each identified fragment can be retained. Furthermore, by acting on intact proteins that retain part of their structure such as disulfide bonds, it is possible to target selected regions for fragmentation such as some of the hypervariable complementarity determining regions (CDRs) that are unique for each clone and necessary for target recognition. Using electron-based fragmentation methods, we employed the prototype timsOmni mass spectrometer to obtain full CDR3 sequences for paired heavy and light chains with high confidence. Optimal results were obtained by performing Electron Induced Dissociation (EID) at ~35 eV electron energy, on native-like fragment antigen-binding (Fab) precursor ions. This approach yields both (a, x) as well as (c, z) fragment ion pairs with the potential to enhance both sequence coverage and annotation confidence. Overall, the timsOmni mass spectrometer presented here serves as an advanced and versatile platform for protein-centric proteomics, demonstrating exceptional sequencing power for comprehensive protein characterization.

## INTRODUCTION

Immunoglobulins, or antibodies, are proteins used by the immune system to target antigens coming from pathogens (such as viruses, bacteria, etc.) or endogenous self-antigens (i.e., auto-antibodies). Matured endogenous antibodies are highly specific to their antigens and can be recombinantly produced and used in therapeutics. Several recombinant monoclonal antibodies (mAbs) are widely used to treat various types of cancer and autoimmune disorders (e.g., rheumatoid arthritis^1^). The approval of antibodies as drugs has increased since their introduction as biotherapeutics in the 1980s; more than 100 mAbs have now been approved by regulatory agencies^2^, most of them IgG1 subtype-like. As antibodies represent a major class of therapeutic agents, their characterization and quality control have become critically important, although these processes remain highly challenging. The structure of the ~150 kDa IgG1s consists of an assembly of two heavy chains (Hc) and two light chains (Lc), held together by multiple disulfide bonds (**Figure 1A**). In the human body, antibodies are produced and secreted by B cells. The amino-acid sequences of these endogenous antibodies’ heavy and light chains are defined by somatic recombination of several V(D)J gene segments.^3^ It has been estimated that humans can potentially produce several billions of distinct antibodies.^4^ Still, a large part of the amino acid sequence of all these antibodies is conserved between all clones (especially in the constant fragment crystallizable (Fc) region of the heavy chains) resulting in an overall high sequence homology (> 80 %),^5^ especially outside of the antigen-binding regions. The antigen specificity of each antibody is defined and constrained in the various hypervariable domains present at the N-terminus of the Hc and Lc, it is therefore important to sequence antibodies to characterize these hypervariable regions and how they relate to the antibodies’ specificity. More specifically, target recognition is driven by three complementarity determining regions (CDR1, CDR2 and CDR3) on each chain. Consequently, sequence coverage of these hypervariable CDR regions, that define the uniqueness of each clone, is of central importance when sequencing antibodies.

**Figure 1.**
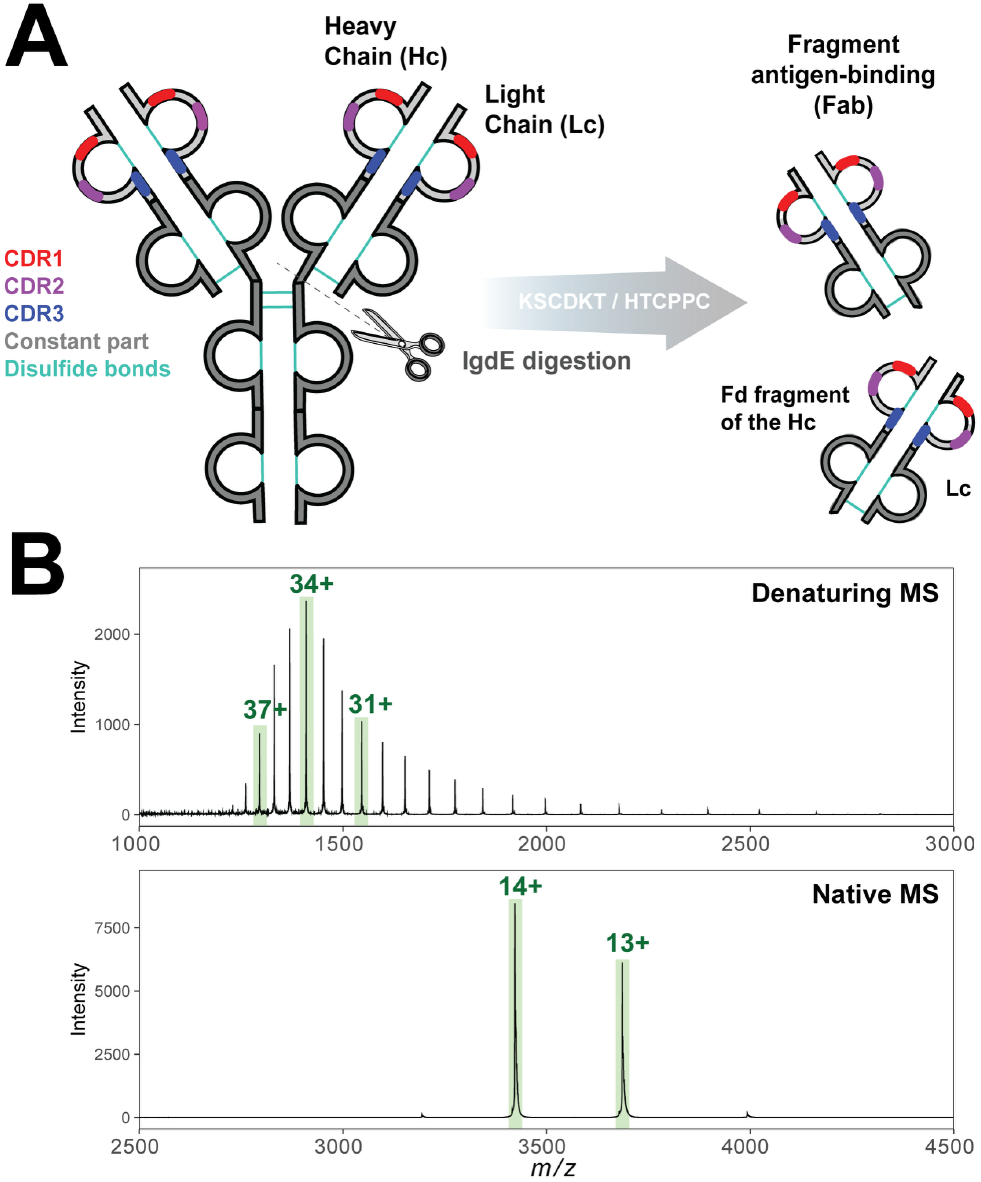
IgG1 Fab-fragment analysis. **(A)** IgG1 antibodies are composed of 2 heavy chains and 2 light chains, each containing 3 distinct and hypervariable complementarity-determining regions (CDRs). IgG1 can be digested by a hinge-cleaving protease, like IgdE, to release the fragment antigen-binding (Fab) regions, reducing the size of the analyte and the impact of the constant Fc part of the antibody on the subsequent analysis. **(B)** the released Fab fragments, which contain still all information on the CDR regions, can be mass analyzed by electrospray under denaturing or native conditions, whereby specific charge states (highlighted in green) can be selected for subsequent tandem MS experiments to obtain sequence information.

Protein sequencing by mass spectrometry (MS) has traditionally been done mostly by using bottom-up (peptide-centric) approaches.^6^ Such methods are also most commonly applied to antibodies, not only for sequencing but also for quality control and multi-attribute monitoring (MAM) in biopharmaceutical analysis.^7^ However, with peptide-centric approaches, the clone as well as the isoform or proteoform origin of the generated/analyzed peptides can be easily lost—a major concern for antibodies, where the pairing of the Lc and Hc and the respective CDRs they harbor, is critical. Protein-centric approaches have therefore emerged as powerful complementary alternatives^8–10^ to study, for example, polyclonal endogenous human antibodies^11–13^ or to monitor biotherapeutics.^14,15^

In recent years, our group has focused on profiling and characterizing the antigen-binding regions of endogenous antibodies from diverse biological fluids.^16–18^ To advance these efforts further, we now describe and deploy a prototype timsOmni platform (**Figure 2A**), thereby extending our work on timsTOF platforms.^19^ Our aim is to ascertain effective fragmentation methods for protein- and proteoform-centric sequencing of precursor ions generated under both denaturing and native MS conditions (**Figure 1B**). To more efficiently access the CDRs at a protein-centric level, an option is to dissect the fragment antigen binding (Fab) domains that contain the most variable parts of the antibody (including all CDRs) from the more sequence-conserved Fc region using hinge-directed proteases, such as IgdE that we employed here. The Fab domain retains the entire Lc as well as a part of the Hc called the Fd fragment, covalently linked through a disulfide bond (**Figure 1A**).

**Figure 2.**
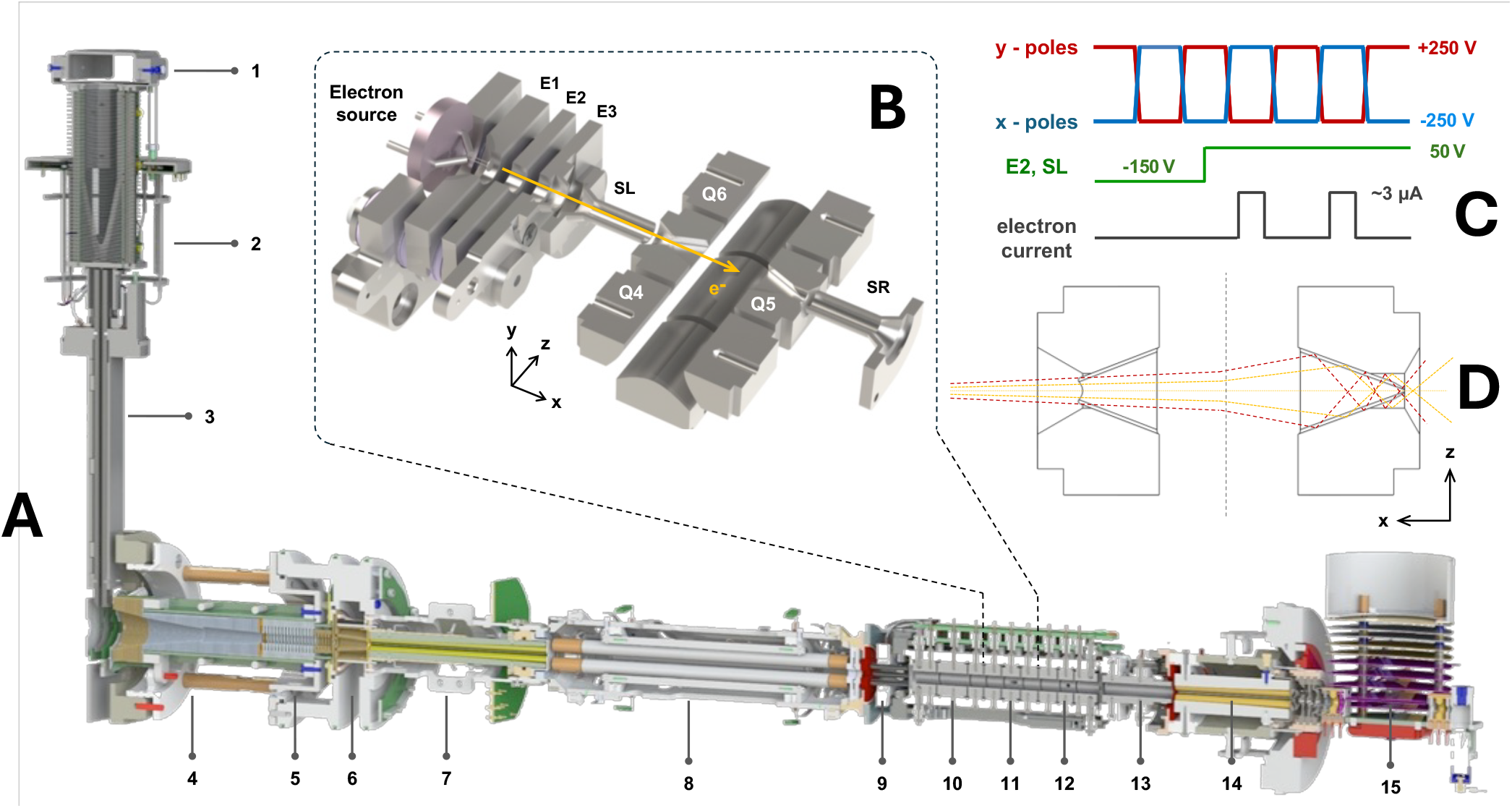
The prototype timsOmni mass spectrometer. **(A)** The prototype timsOmni instrument is based on a timsTOF Ultra and incorporates the latest revision of the Omnitrap platform. Key components are labelled and include: (1) ion inlet, (2) intermediate pressure ion funnel, (3) RF quadrupole ion guide, (4) accumulation and (5) mobility analyzer regions of the trapped ion mobility spectrometer, (6) low pressure ion funnel, (7) RF quadrupole ion guide with axial DC field, (8) quadrupole mass filter, (9) RF hexapole ion guide, the Omnitrap platform including (10) section Q2 configured for resonance excitation CID and resolving DC isolation, (11) section Q5 enabling ExD ion processing, resonance excitation CID and resolving DC isolation, (12) section Q8 providing optical access and (13) new section Q10 for ion transfer, (14) collision cell and (15) the orthogonal acceleration region for sampling ions into a reflectron TOF mass analyzer. **(B)** Cross sectional view of the external electron source comprising a hot disc cathode and a system of lenses externally coupled to section Q5. A dense electron beam is collimated into the trapping region enabling processing of ions via ExD with fine control of the electron energy ranging from 0 to 150 eV. **(C)** Two antiphase rectangular waveforms are applied to the Omnitrap pole-electrodes while electron injection in Q5 is accomplished during the positive phase of the rectangular waveform applied to the X-poles and further enabled by a switched DC signal applied to a lens-electrode. **(D)** Cross sectional view of section Q5 on the XZ plane illustrating the newly implemented Q5 inlet and outlet slot design for enhanced electron transmission and durability, including a schematic representation of higher-energy electron trajectories within a static quadrupole field.

For protein-centric characterization of intact proteins and protein complexes fragmentation methods alternative to collision-induced dissociation (CID) have been shown to be beneficial.^9,20^ These include several laser-based approaches, for example, ultraviolet photodissociation (UVPD)^21,22^ and Electron-based Dissociation (ExD) methods, among which the most commonly reported are Electron-Transfer Dissociation^23,24^ (ETD) and Electron-Capture Dissociation^25,26^ (ECD). While ETD involves the reduction of a polycation by a radical reagent anion, ECD entails the capture by a polycation of one or more low-kinetic-energy free electrons. Both result in the formation of intermediate highly energetic charge-reduced radical species (**Equation 1**), that can subsequently dissociate via bond cleavage.^27,28^

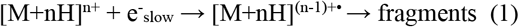

Illustrative pioneering data obtained with ECD and ETD for the characterization of antibodies started with the analysis of intact mAbs as early as the 2010s,^29,30^ further expanded more recently with data on Lc/Hc pairing^31^ and the protein-centric sequencing of relevant CDR sequence tags,^31,32^ even with precursor ions up to MDa IgM complexes.^33^ ECD/ETD mostly generate sequence ladders of c and z ions, but have been successfully combined with other non-collisional activation approaches such as UVPD and infrared multiple photon dissociation (IRMPD) for improved sequence coverage, including better coverage of the CDR regions.^31,34^ Knowing that UVPD can produce different types of fragments compared to ECD, notably (a, x) ion pairs,^35–37^ it is worth exploring activation methods that could produce correlated ion pairs in a single analysis. Here, we specifically explore whether higher-electron-energy methods, e.g., Electron-Induced Dissociation (EID), could be employed for such purposes. The term EID serves as an umbrella designation for electron-based fragmentation techniques that employ electron energies above ~10 eV. These methods induce ion fragmentation through electronic excitation and/or ionization, thereby providing access to additional dissociation pathways beyond those available through ECD. Here, we apply EID with 35 eV electrons (**Equation 2**), where the slow electrons released during the process may be recaptured (**Equation 1**).^38^

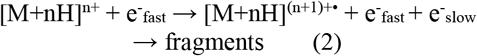

We first describe and evaluate a newly designed prototype platform integrating an Omnitrap linear ion trap into a timsTOF-based instrument and employ it as a proof-of-concept for the characterization and sequence analysis of antibodies. We provide a detailed overview of the newly designed Q5 in the Omnitrap platform that allows us to perform low-energy ECD and high-energy EID on precursor ions of Fabs generated by electrospray ionization under either denaturing or native MS conditions. The specific aim was to obtain unambiguous annotations of the CDRs of the antibodies, ideally with correlated reads of several fragment ion series (i.e., (c, z), (a, x)), each present in multiple charge states, which would increase confidence in sequence assignments from *de novo* sequencing of hypervariable regions of antibodies.

## RESULTS

### timsOmni prototype instrument design and description

This work presents the first out-of-factory application of a prototype timsOmni platform (Bruker), used here to characterize and sequence Fabs from IgG molecules, utilizing and comparing both low energy ECD and high energy EID techniques. The hardware combines an extensively modified timsTOF Ultra geometry with the latest Omnitrap technology. The Omnitrap platform can function in trapping mode, with ExD and CID fragmentations, implementing intermediate ion enrichment steps enabling deep sequencing through highly efficient tandem MS^n^ workflows, or in transmission mode, maintaining all data-independent acquisition (DIA) flavors combined with parallel accumulation serial fragmentation (PASEF) available on timsTOF mass spectrometers.

The configuration of the prototype instrument excluding the time-of-flight mass analyzer is depicted in **Figure 2A**. Ions generated via electrospray ionization enter the system through a 1.0 mm inner diameter resistive glass capillary (1) into the first fore-vacuum chamber, which is maintained at 10 mbar pressure. Within this chamber, ions carried by the under-expanded jet are redirected at a right angle to the jet axis by a deflector lens, collected by a radio frequency (RF) ion funnel (2) and focused through a differential aperture into an RF quadrupole ion guide (3). The quadrupole ion guide is enclosed within an insulating jacket, establishing laminar flow conditions and facilitating effective ion transport through the secondary fore-vacuum stage of the instrument. This design eliminates the necessity for an additional axial direct current (DC) field component to drive ions downstream. A second deflector lens is employed to redirect ions into a high-capacity trapped ion mobility spectrometer (TIMS) configured with an accumulation region (4) and an ion mobility analyzer region (5) operated at 2 mbar pressure. Ions transported through the TIMS device can be accelerated towards a second RF ion funnel (6) maintained at 0.2 mbar to a maximum kinetic energy of 200 eV to enhance desolvation. Accelerated ions are re-thermalized at 0.01 mbar pressure in a second RF quadrupole ion guide (7) designed with a suitable set of auxiliary electrodes forming an axial electric field to deliver a collimated beam of ions into downstream optics. Mass selection is performed using a quadrupole mass filter (8) equipped with a modified RF generator extending the precursor ion isolation window to 4500 Th.

The Omnitrap platform is situated downstream of the quadrupole mass filter. Efficient coupling between these elements is achieved through a short RF hexapole (9) and a series of DC lenses. The segmented architecture of the Omnitrap platform allows ions to be directly accumulated and processed within multiple designated regions. Sections Q2 (10) and Q5 (11) facilitate CID via resonance excitation and ion isolation using resolving DC signal components, while section Q5 (11) is further configured to receive a pulsed electron beam from an external electron source for ExD fragmentation of trapped ions. Section Q8 (12) provides optical access for photodissociation and can also serve as a storage region for product ions to enhance signal intensity. Additionally, quadrupole segment Q10 (13) has been incorporated to guide ions from the Omnitrap platform to the collision cell (14), which is configured with a gate lens to release ions into the gridless orthogonal acceleration region (15) of a reflectron TOF mass analyzer.

**Figure 2B** illustrates the newly implemented section Q5. An yttria-coated iridium disc cathode with 1.6 mm diameter (Kimball Physics) is driven to incandescence temperatures (~4.6 A heating current) producing >20 μA of electron current. A fraction of this current collimated through a system of high-voltage lenses (E0 - E3, SL) is injected into the trapping region where ions are radially and axially confined by the two antiphase rectangular waveforms (±250 V_0p_, 0.1-2.4 MHz) and appropriate DC offset potentials applied to sections Q4 and Q6, respectively. Electron energy is finely tuned by adjusting the voltage difference between the DC bias applied to the cathode and the DC offset applied to section Q5. The energy spread of electrons available in ExD reactions is dictated primarily by the number of charges stored in Q5 and the corresponding radial size of the trapped ion cloud (<5 eV under extreme space conditions). Electrons are injected into the trapping region of Q5 only during the positive phase of the rectangular waveform applied to the X pole-electrodes of the Omnitrap platform (**Figure 2C**). The electron current accessible for ExD reactions is estimated at approximately 3 μA. Switched DC pulse signals applied to lens electrodes E2 and SL prevent stray electrons from entering the trapping region. The deflected electron beam is collected by a positively biased conductive casing enclosing the electron source. To mitigate surface charging and eliminate unwanted effects during extended ion trapping, all proximal ceramic components are covered with grounded conductive masks.

Further design and performance improvements compared to the previously published geometry^39^ were realized, considerably improving robustness and reaction speed by carefully shaping the inlet and outlet openings on the X pole-electrodes of the Omnitrap platform (**Figure 2D**). The XZ cross-sectional view of section Q5 illustrates the new inlet design comprising a tapered entrance leading to a restriction interface to reject off-axis electrons, followed by a divergent two-dimensional channel terminating to a 2×8 mm^2^ exit aperture to minimize electron reflection from surfaces. An asymmetric configuration is adopted on the outlet X pole-electrode accounting for (i) the quadrupole field symmetry field required to confine ions efficiently for hundreds of milliseconds, (ii) the electron defocusing that occurs when traversing a static quadrupole field, and (iii) the reflection of electrons from surfaces and their subsequent guidance through the outlet toward a positively-biased collector. This configuration effectively eliminates surface contamination on the hyperbolic pole-electrodes of section Q5 and suppresses fast stray electrons that would otherwise react with trapped ions, evidenced by the absence of b- and x-type fragments, which were observed in the initial ECD experiments with proteins.^39^ The redesigned system thus establishes pure ECD conditions, yielding primarily (c, z) ion pairs.

Ions selected by the quadrupole mass filter according to their *m/z* values are directly accumulated in section Q5 for ExD fragmentation. Experimental measurements indicate that section Q5 has a capacity of approximately 50 million charges for isolated charge states of multiply charged proteins. At an incoming ion current of 1 nA (equivalent to 6.24 million charges per ms), this capacity is reached in approximately 8 ms. For lower abundance species, accumulation times may be extended to 500 ms or longer to obtain ExD mass spectra with adequate signal-to-noise ratios suitable for confident annotations.

The orthogonal acceleration time-of-flight (OA TOF) mass analyzer operates at a 5 kHz repetition rate (200 μs flight time), enabling efficient sampling of ions generated either directly from the electrospray ionization (ESI) source or in the form of product ion pulses ejected from the Omnitrap platform. For example, a 10 ms product ion pulse ejected from section Q5 containing 10 million charges produces 50 TOF extraction events, each sampling approximately 200,000 charges. The high repetition rate of the timsOmni platform accommodates such ion fluxes, delivering high sensitivity and signal-to-noise ratio mass spectra at high acquisition speeds. The standard OA configuration, when paired with a short-flight time TOF mass analyzer, offers better handling of high ion flux compared with direct ejection to TOF from trapping devices (where space charge effects in RF fields can compromise ejection efficiency, transmission, and TOF resolving power) or with multipass TOF systems (which operate at reduced repetition rates due to multi-millisecond long flight times).

### Assessing the performance of the prototype timsOmni platform for protein-centric characterization of antibodies

We hypothesized that some key features of the new prototype timsOmni described here would be very beneficiary for top-down sequencing of especially the hyper-variable segments of antibodies. To support this, we aimed here to extend upon the work of Shaw^31^ and Greisch^32,33^, who demonstrated that electron-induced fragmentation of antibodies—or their F(ab’)2, Fab fragments—yields unique insights into key regions of the antibody sequence, particularly the CDR regions critical for antigen binding, with an emphasis on the key CDR3 segments. We especially wanted to explore how different electron energies, as available on the timsOmni, would affect the observed sequence information, and whether this would be optimal when analyzing highly charged denatured Fab molecules (like those generated by standard LC-MS), or their relatively low charged variants generated by native electrospray. We focused within this proof-of-concept work primarily on the Fab fragments of two recombinant monoclonal antibodies, namely trastuzumab-IgG1 and 7D8-IgG1 (the former kindly provided by Roche, the latter by Genmab).

**Figure 3** shows representative electron energy dependence curves obtained for individual charge states of native Fab species, illustrating how electron energy governs the formation yields of the charge-reduced and meta-ionized species in ECD and EID experiments, respectively. Indeed, looking at the charge-reduced or charge-increased precursors is the most straightforward way to monitor the efficiency of the ExD processes. In ECD, two optima are identified by monitoring the intensity of the 13+• charge state normalized to the 14+ precursor ion. The maximum observed at approximately 0 eV corresponds to the optimum electron energy for ECD, whereas the shoulder at 7 eV reflects the broader energy window characteristic of hot ECD. Notably, the hot ECD tail extends substantially beyond the range previously reported for smaller proteins such as insulin and ubiquitin.^39^ The ECD response curves for multiply charged Fabs can in part be attributed to an electron recapturing process, in which either the ionizing electron or a detached low energy electron is attracted back to the protonated ion, creating another radical site and inducing electronic excitation.^38^ In addition, electronic-to-vibrational energy transfer processes on the millisecond time scale contribute to the diverse fragment types observed in EID experiments with electron energies exceeding the ionization threshold (>10 eV). Finally, space charge can further influence the energy dependence by controlling the ion cloud size and thereby influencing the energy distribution of electrons driving the ExD, (x = C, I) reactions. In contrast to ECD, the energy dependence in EID appears largely insensitive to protein size (~50 kDa) and precursor charge state (z<20+) within the range investigated, with the maximum ionization efficiency attained at 35 eV electron energy.

**Figure 3.**
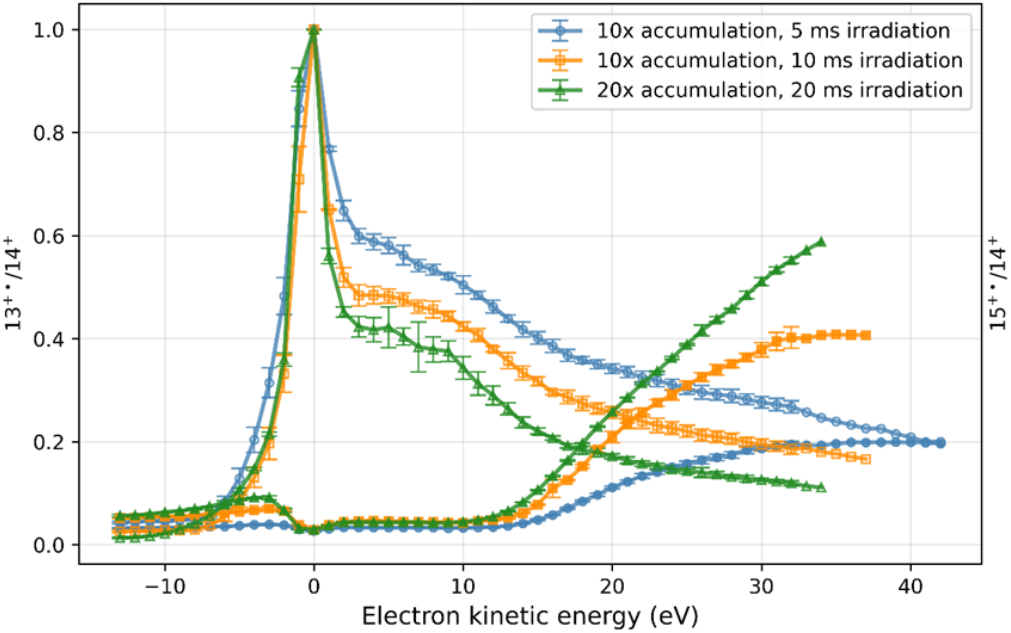
Electron energy dependence curves monitoring the efficiency of electron capture. We monitored the electron capture 13+•/14+ intensity ratio (open symbols) and the meta-ionization 15+•/14+ (filled symbols) processes for the 14+ charge state of the 7D8-IgG1 Fab ionized under native conditions. The blue circles, orange square, and green triangle symbols correspond to an accumulation and irradiation of (100 ms, 5 ms), (100 ms, 10 ms), and (200 ms, 20 ms), respectively. For convenience, the data sets were normalized relatively to the maximum of the 13+•/14+ curve at near zero electron kinetic energy.

#### ExD affords critical insights into the Hc-Lc pairing and sequences of Fab molecules

Having established the optimal electron energies for ECD and EID, we next aimed at benchmarking the fragmentation of IgG1 Fab molecules to identify the strengths and weaknesses of low-electron-energy ECD and higher-electron-energy EID. We analyzed Fabs ionized under denaturing conditions, representing how they are typically generated in a liquid chromatography setup coupled to MS (LC-MS), and also under native conditions. The most intense charge state of the Fab in the MS1 mass spectra (**Figure 1B**) was selected for ExD fragmentation and subjected to electron irradiation for 50 ms (if denatured) or 100 ms (if native). As expected, the ExD-MS/MS spectra of the 34+ and 14+ precursors (**Figure 4** for 7D8-IgG1, **Figure S1** for trastuzumab) showed prominent charge-reduced Fab molecules in ECD whereas EID mainly produced charge-increased precursor ions.

**Figure 4.**
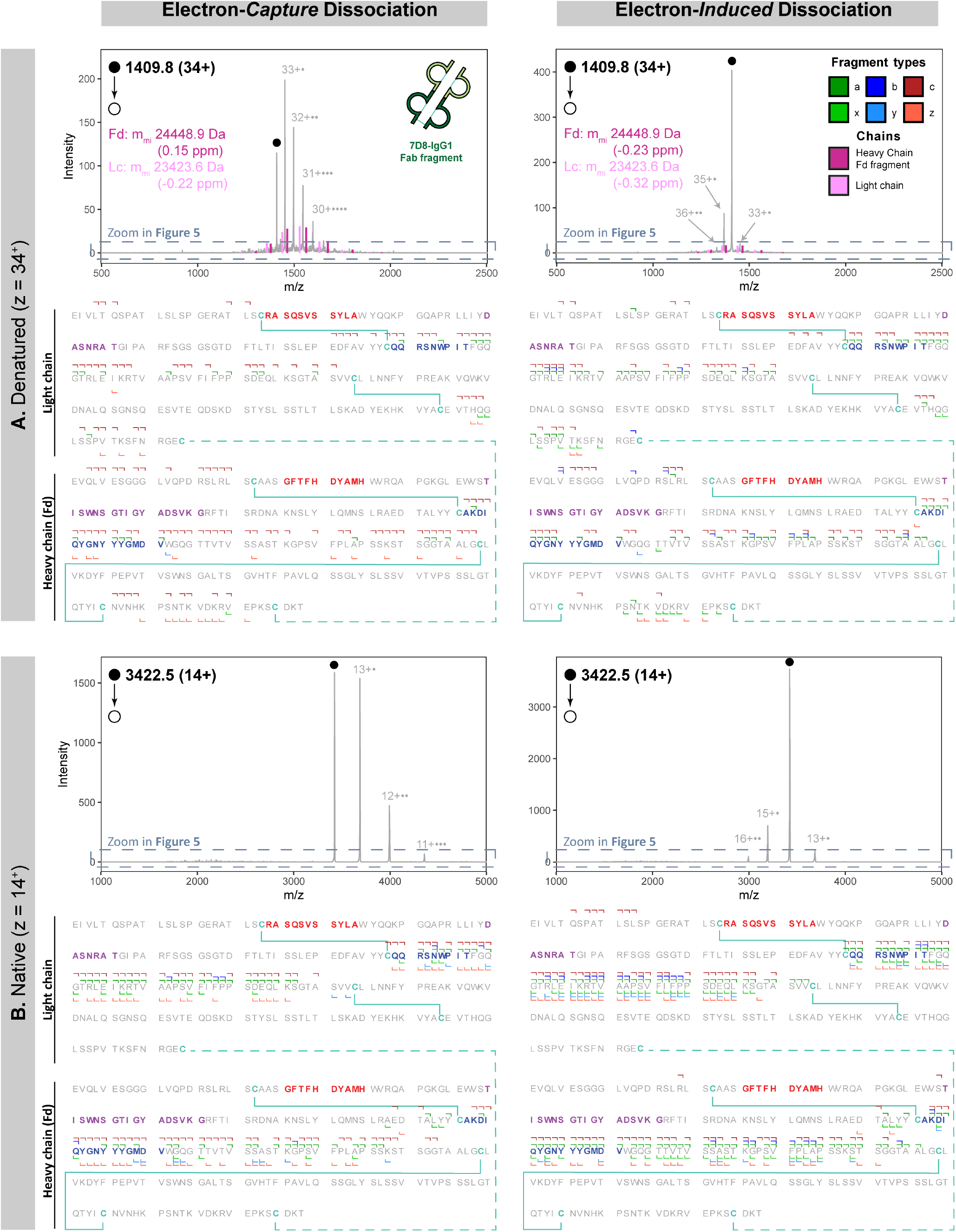
ExD provides information on Lc/Fd pairing and delivers unambiguous sequence coverage of the CDR3 region through multiple complementary fragment ion series. ExD-MS/MS spectra of the 7D8-IgG1 Fab with confidently annotated fragment ions mapped onto its sequence, highlighting CDR1, CDR2, and CDR3 in red, purple, and blue, respectively, for **(A)** higher charge state Fab ions produced under denaturing conditions (z = 34^+^) and **(B)** lower charge state Fab ions produced under native conditions (z = 14^+^), each analyzed by low-energy ECD (left) and high-energy EID (right).

Importantly, for the denatured Fabs, the next most intense signals corresponded to the Lc and the Fd fragment of the antibodies, observed in both ECD and EID spectra across at least five charge states each, thus enabling their identification with sub-ppm average accuracy. The data reveals that under the selected operating conditions, ExD methods on the timsOmni can promote gas-phase reduction of the interchain disulfide bond, leading to the release of both Lc and Fd chains (**Figure 4A**). These fragmentation schemes are in line with earlier findings^31^ and are key to retrieve information about Lc/Fd pairing of antibodies. In contrast, in the case of native Fab molecules, Lc and Fd fragment ions were not observed in either ECD and EID experiments, presumably due to the non-covalent interactions stabilizing the two chains and since no supplementary collisional activation was applied in these experiments to dissociate the radical Fab ions.

Under denaturing conditions, adequate sequence coverage was achieved by ExD for the most easily accessible regions on both chains, (i.e., the N- and C-terminus, and the regions between the two intrachain disulfide bonds, **Figure 4A)**. The sequence of the CDR3 stretch, being in the region between the intrachain disulfide bonds, could already be deciphered using ECD by a sequence read of c-type fragment ions (90 % coverage for the Lc-CDR3, 88 % coverage for the Hc-CDR3). With EID, a correlated series of a-ions provided even greater confidence in CDR3 sequence determination, reaching 100 % coverage for Lc-CDR3 and a slightly lower coverage of 75 % for Hc-CDR3. Alike results were observed when performing ECD and EID on other denatured precursor charge states (**Figures S2**, 31+ and 37+), whereby sequence information was largely independent of the precursor charge state. CDR1 and CDR2 could not be directly covered by MS2 ExD as they are enclosed within an intrachain disulfide bridged loop, in line with what has been observed and noted earlier.^31–33^

Under native conditions, ExD provided excellent coverage of the region in-between the intrachain disulfide bonds, encompassing the CDR3 (**Figure 4B**). Already by using ECD, most of this region was identified by at least three different fragment series (a, c and z) yielding a sequence coverage of 100 % for the Lc-CDR3 and 88% for the Hc-CDR3. Even better, with EID, the sequence exhibited the same coverage of 100 % for the Lc-CDR3 and 88 % for the Hc-CDR3, but this time with coverage by four full sequence series of a, c, x and z ions, occasionally supplemented by a few b and y ions. Under native conditions the fragments originating from the Lc had slightly higher intensity than those of the Fd fragment of the Hc (**Figure S3**), explaining why there are some gaps in the C-terminal series of the Fd compared to the Lc. In contrast to denatured Fab molecules, the charge states of the native precursor ions appear to impact the observed fragments, especially on the C-terminal ions (**Figures S4A**, 13+), a phenomenon which will be discussed further below.

Similar results were obtained for the Fab of Trastuzumab (**Figure S1**) as for the 7D8-IgG1 Fab described above. Overall, under native conditions, EID outperformed ECD in providing unambiguous sequence information for the hypervariable CDR3 region through the parallel detection of multiple sequence reads. Interestingly, the opposite trend was observed for denatured Fab precursor ions where ECD provided better CDR3 coverage compared to EID. We hypothesized that this difference could be linked to spectral congestion caused by extensive overlap between radical precursor and the fragment ion distributions, complicating confident assignments.

To further investigate the difference in sequence coverage observed between denaturing and native conditions, we compared the distributions of the various fragment ion types across the *m/z* range (**Figure 5**).

**Figure 5.**
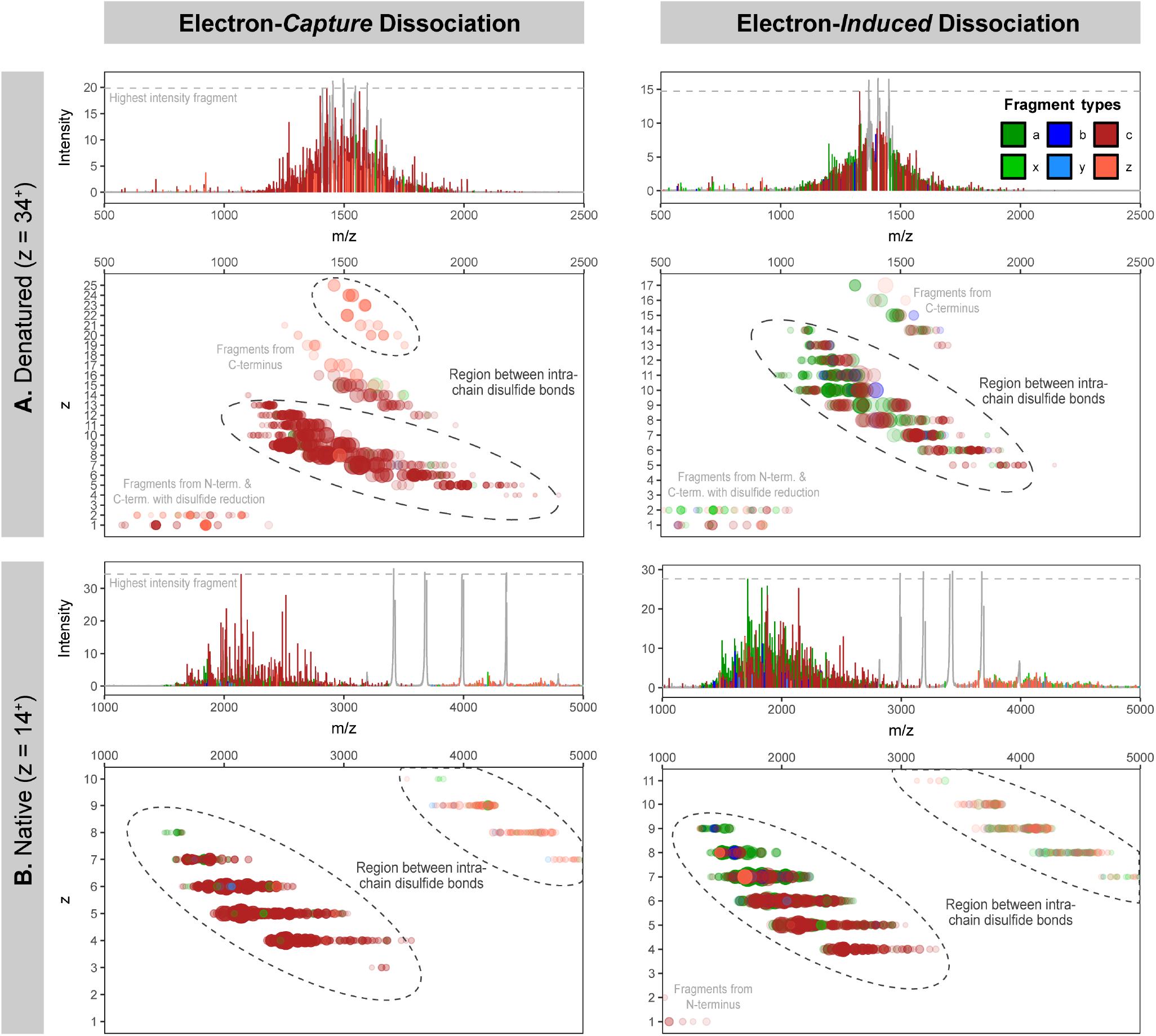
Improved separation across the m/z range results in more interpretable data when Fabs are analyzed under native MS conditions. Low-energy Electron-Capture Induced (ECD, left) and high-energy Electron-Induced Dissociation (EID, right) MS2 spectra (as in **Figure 4**, but with an expanded y-axis to better visualize lower-intensity, color-coded fragment ions) are shown for **(A)** the 34+ charge state of 7D8-IgG1 and **(B)** the 14+ charge state of 7D8-IgG1. The corresponding charge state versus m/z plots of the annotated fragment ions (dot size reflecting intensity and transparency indicating confidence) demonstrate that, under denaturing conditions, multiple fragment ion series from different regions of the chains (i.e., high- and low-mass fragments) overlap extensively in the m/z dimension—the distributions relevant to CDR3 characterization are circled and annotated in black, other distributions are annotated in gray. In contrast, when precursor ions are generated under native conditions, these ions become distinctly separated, enabling more accurate spectral interpretation.

When highly charged Fab precursor ions are produced under denaturing conditions, nearly all fragment ions produced in ECD and EID are located in the 1000-2000 Th range, overlapping with the radical precursor ions formed by electron capture and electron ionization, respectively. The spectral complexity is increased by the formation of multiple fragment ions observed at different charge states corresponding to the same cleavage position (**Figure 5A**). In ECD, most fragments originate from the regions between the two intrachain disulfide bonds on both chains. The majority of the C-terminal fragments also retain the second chain (Lc or Fd) via the interchain disulfide bond. Additionally, ECD fragments from the constant domains of both chains near the C-terminus (intermediate charge states) are observed in the same *m/z* region, further increasing spectral complexity. In EID, fragment ions are produced in similar regions but with higher charge states, reflecting their origin from ionized Fab precursor species. These fragmentation features cause significant overlap among fragment ions (x-axis zoom in **Figure S5**), as evidenced by the low number of z-ions in ECD and their low-confidence assignment (shown by their high-level transparency in **Figure 5A**).

Fragment ions from both chains are observed at comparable abundance across the two ExD methods (**Figure S3**). Specifically, ECD predominantly yields c-type fragments along with lower-intensity a- and z-type ions, whereas EID produces a more balanced distribution of a- and c-type ions, accompanied by low-abundance b-ions (**Figure S6A**). Based on the data obtained on Trastuzumab- and 7D8-Fabs analyzed under denaturing conditions, EID demonstrated a greater potential for delivering unambiguous sequence reads compared to ECD, primarily due to the additional formation of the a-ion series. However, the substantial spectral complexity still constrains annotation quality and ultimately limits overall sequence coverage.

When lower charge state Fab ions are generated under native conditions (**Figure 5B**) and subjected to ExD activation, similar fragment ions are observed; however, they are distributed across a much broader *m/z* range (1000–5000 Th). Two main fragment populations are evident in both ECD and EID, originating from the region between the intrachain disulfide bonds. The first, and most intense, population appears at lower charge states between 1500 and 3500 Th and consists primarily of N-terminal fragments—predominantly c-type ions in ECD and a- and c-type ions in EID (**Figure S6B**). The second population, extending from 3500 to 5000 Th, corresponds to complementary C-terminal fragments that retain the counterpart chain, mainly z-type ions in ECD and x- and z-type ions in EID. Under native MS conditions, the precursor charge state notably influences the distribution and intensity of these C-terminal fragment ions (**Figure S4B**).

From these results, the advantages of generating lower charged Fab precursor ions under native MS conditions are evident: (i) the resulting complementary populations of fragment ions are well-resolved in the *m/z* dimension, and separated from the Fab precursor and its related radical ions, (ii) the reduced charge states of the fragments yield better-resolved isotopic patterns, facilitating more confident identification (**Figure S7**), and (iii) the intermediate C-terminal fragment population observed under denaturing conditions (**Figure 5A**) is not observed, thereby avoiding interference from ions outside the region of interest. It is noteworthy that high-mass x- and z-type ions were identified with lower confidence than a- and c-type ions. This is partly due to their partial isotopic resolution (**Figure S8**), and also to suboptimal transmission of high-mass fragments in this first-generation timsOmni prototype, resulting from ion-optical limitations between the collision cell and the TOF analyzer. Ongoing developments in the next-generation timsOmni design have addressed these transmission constraints.

#### ExD provides high-confidence CDR3 annotations across the region encompassing CDR3

To further assess the influence of electron energy on ExD efficiency and its dependence on the Fab precursor charge state, we evaluated annotation quality using the OmniScape data analysis software, applying two distinct scoring metrics (**Figure 6**). (i) The annotated-fragment confidence score indicates how closely the theoretical isotopic distributions match the experimental patterns, reflecting the quality of individual annotations; (ii) The residue confidence score (depicted in blue in **Figure 6**), calculated for each amino acid separately, represents the overall confidence for all fragments bracketing a given residue (see details in the **Supplementary Information**).

**Figure 6.**
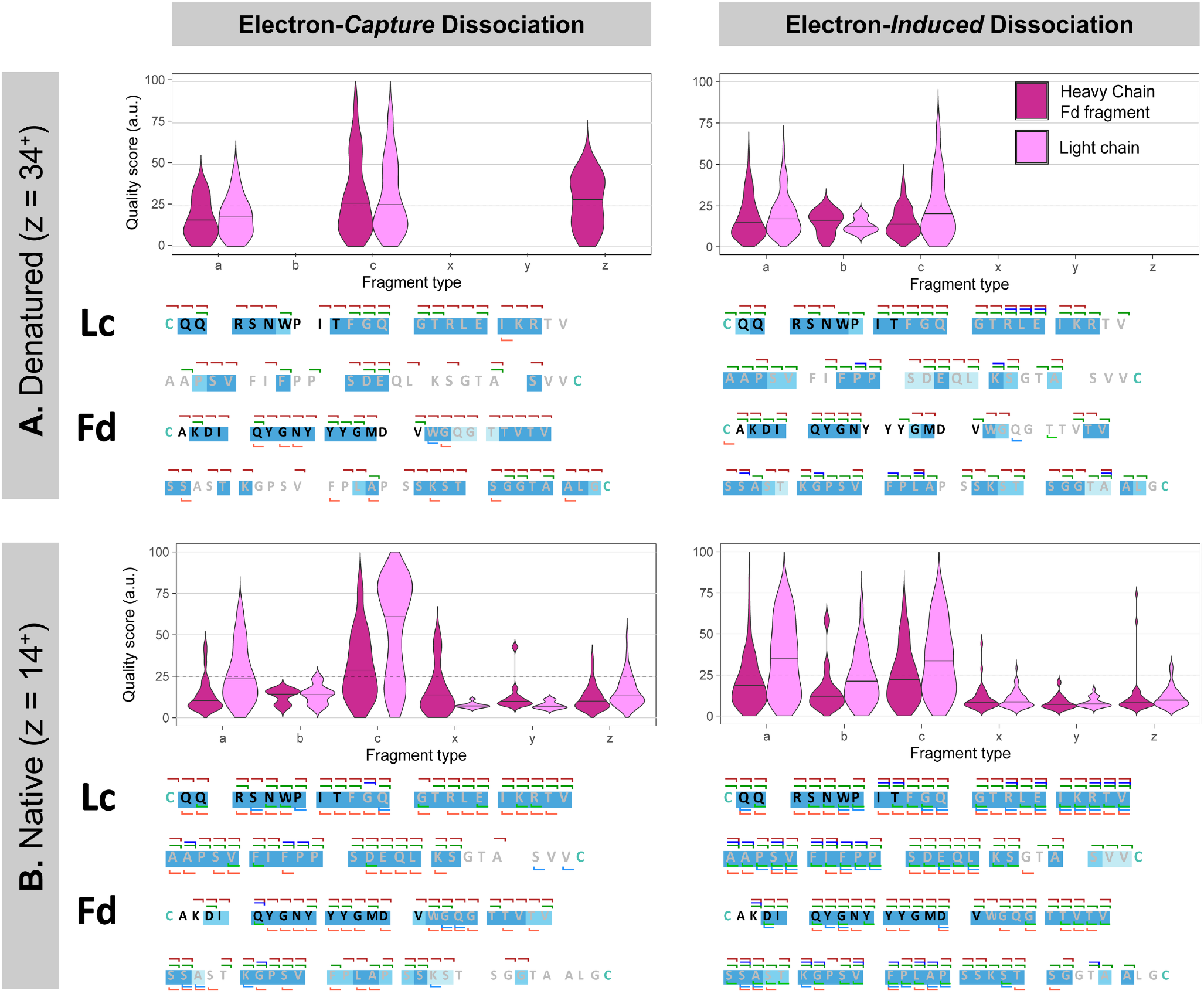
Electron-Induced Dissociation at 35 eV of native-like Fab precursor ions provides the most confident annotation of the hyper-variable regions of immunoglobulins. In this analysis, only ions corresponding to the regions between the two intrachain disulfide bonds— encompassing the CDR3 (highlighted in black)—were considered. The violin plots illustrate the distribution of the annotated-fragment confidence scores for each fragment ion type identified in the ExD MS^2^ spectra (solid line: median; dashed line: high-quality threshold). Below each violin plot, residue confidence scores are mapped onto the sequences, with the darkest blue indicating the amino acid annotations with the highest level of confidence. The left panels show results obtained by low-electron-energy ECD, while the right panels display those produced by higher-electron-energy EID for **(A)** the higher charge state precursor ions produced under denaturing conditions (e.g., 34^+^ 7D8-IgG1 Fab) and **(B)** lower charge state precursor ions produced under native conditions (e.g., 14^+^ 7D8-IgG1 Fab). Overall, high-energy EID of native-like precursor ions provides the most comprehensive sequence coverage of the immunoglobulin hypervariable regions, whereas ECD produces fragments with the highest individual quality scores under both native and denaturing conditions.

First, the annotated-fragment confidence score was calculated for fragment ions originating from the regions between the intrachain disulfide bonds (7.5–17.5 kDa and 30– 40 kDa), which are relevant for the characterization of the hypervariable CDR3. This score ranges from 0 to 100, with 100 indicating the highest confidence in fragment annotation. In practice, scores above 25 (dashed lines) were found to correspond to high-confidence annotations.

Higher confidence scores were observed for the c-fragment ions compared to their corresponding a-type counter-parts in ECD experiments with the higher charge state Fab precursors generated under denaturing conditions. In contrast, EID produced lower scoring c-ions but yielded more complete series of additional a- and b-type fragments (**Figure 6A**). Under native conditions, ECD produced fragments with increased confidence scores, particularly for the c-type ions originating from the Lc. The most comprehensive sequence coverage was obtained in native EID mode providing high-confidence a-, b- and c-type fragment annotations from both chains, extending across the entire region of interest (**Figure 6B**).

Overall, considering the high confidence of all N-terminal fragment series and the presence of the complementary C-terminal fragments, high-energy EID of native-like precursor ions provided the most reliable annotation of the immunoglobulin hypervariable CDR3 regions. Nevertheless, ECD offers complementary advantages, most notably generating higher-quality scores and unique insights into the Lc/Fd pairing, as described above.

## CONCLUSIONS

A prototype timsOmni MS mass spectrometry platform is described, featuring the latest iteration of Omnitrap technology fully integrated into the well-established timsTOF Ultra platform. We evaluate the performance of an improved Q5 section within the Omnitrap platform dedicated to ExD, which offers enhanced durability and accommodates higher electron currents. This design enables faster electron-based activation schemes compared to the original configuration, facilitating the acquisition of high-quality ECD and EID spectra.

In this proof-of-concept study, we employed the instrument to sequence IgG1 Fab fragments, building on previous findings that demonstrated the utility of ExD for obtaining sequence information from CDR3 variable regions. In this context, we evaluated the potential of the timsOmni platform using both higher and lower charge state Fab molecules produced under denaturing and native conditions, respectively. Fragmentation behavior was compared between low-electron-energy ECD and higher-electron-energy EID.

Focusing on the CDR3 region of the Fab molecules, we evidenced that the best coverage and highest confidence were reached by deploying EID on native-like lower charge state Fab precursors. This approach produced multiple correlated sequence reads and highly confident N-terminal fragment ion assignments, largely due to reduced spectral congestion. In contrast, ECD additionally yielded high-intensity Lc/Fd fragment ions informative for chain pairing, along with slightly more high-confidence c-ion reads. These features were particularly prominent for Fab species generated under denaturing conditions. Since chromatographic separation of Fabs is generally done using reverse-phase LC-MS that yields denatured Fabs,^19^ this suggests that ECD is better suited for reverse-phase LC-MS workflows and could be used for online Lc/Fd pairs determination.

The observed recurring fragments arising from identical cleavage sites contribute substantially to spectral congestion, however, the presence of correlated fragment types across multiple charge states represents a valuable spectral characteristic for advancing *de novo* sequencing algorithms. Such redundant fragmentation can help minimize false-positive identifications and enhance the confidence of *de novo* sequence tags derived from complex mass spectra.

Although CDRs 1 and 2 are not directly accessible by MS2 ExD due to the presence of intrachain disulfide bridges, these regions could become accessible by applying supplemental collisional activation to the ExD product ions or via the development of higher-order MS^*n*^ workflows, which can be readily implemented on the Omnitrap platform. While the development of comprehensive protein-centric MS^*n*^ strategies remains ongoing, the MS2 ExD approach described here is already compatible with liquid chromatography workflows, enabling the characterization of biotherapeutics and even polyclonal biological samples, as will be detailed in a forthcoming publication. Continuing efforts to improve mass spectrometry-based technology are expected to advance the applicability of protein-centric native LC-MS approaches.^40^

In this proof-of-concept study, we focused on the top-down characterization of antibodies using ECD and EID, with the intention of extending our work to include supplemental collisional activation of ExD product ions and laser-induced photodissociation techniques such as UVPD and/or IRMPD, thereby demonstrating the versatility of the timsOmni platform. With these fragmentation methods available, diverse experimental schemes can be developed to enable peptide- and protein-centric MS^n^ strategies, supporting the analysis of a wide range of biomolecules, including histones, glycopeptides, glycoproteins, phosphoproteins, microproteins, and even intact protein complexes. The capabilities of the timsOmni mass spectrometer will be further enhanced through the implementation of MS^n^ strategies combined with ion mobility, enabling new analytical workflows in the next-generation timsOmni design.

## MATERIALS AND METHODS

### Fab preparation

Fab fragments were produced from two recombinant monoclonal IgG1 antibodies, Trastuzumab (Roche, Penzberg, Germany) and 7D8 (produced by Genmab, NL). These antibodies were digested by incubating 200 µg of each antibody with the IgG1-specific hinge-cleaving protease immunoglobulin degrading enzyme (IgdE) overnight (> 16 hours), with a 10:1 (mass/mass) antibody to enzyme ratio at 37 °C in 150 mM phosphate buffer, pH 7 (50:50 monosodium and disodium phosphate), under agitation at 750 rpm. Fab fragments were then purified by (i) removing the Fc fragments by incubating with 50 µL of CaptureSelect FcXL (Thermo Fisher Scientific) slurry for 1 hour at room temperature with agitation (750 rpm), then (ii) removing the His-tagged IgdE by incubating with 50 µL of Ni-NTA beads slurry for 30 min at room temperature with agitation (750 rpm). The reference Fabs were then prepared for analysis under two conditions. The first half was prepared for native MS, and directly buffer-exchanged into 150 mM ammonium acetate using 7K Zeba spin columns (Thermo Fisher Scientific) (2 cycles). The other half was prepared for denaturing MS and first denatured using 8 M guanidine in a 1:3 ratio and incubated for 15 min at 60 °C (650 rpm), then buffer-exchanged in 20 % acetonitrile + 0.1 % formic acid using 10 kDa molecular weight cut-off Amicon filters (Merck Millipore) (7 cycles).

### Mass spectrometry

For the proof-of-concept measurements reported here, a prototype timsOmni mass spectrometer was equipped with an in-house-fabricated 3D-printed ionization source (**Supplementary Figure S9**), enabling static nano-electrospray ionization (nESI) to be performed under native or denatured conditions using gold-plated borosilicate emitters.

#### ExD fragmentation of denatured Fab precursor ions

Denatured Fabs were diluted in 80:20 water/acetonitrile with 0.1 % (v/v) formic acid to a concentration of 5 μM and loaded in aliquots of ~ 3 μL in in-house pulled emitters for nESI. All ions were accumulated for 30 ms in the TIMS cartridge (the maximum value before the onset of space-charge effects, which may lead to ion losses), submitted to an additional collisional activation (in-source CID) of 40 eV, and then charge states 31+, 34+, and 37+ were mass isolated using the quadrupole mass filter with an isolation width of 5 Th (see **Supplementary Table S1** for detailed instrument parameters and **Supplementary Table S2** for precursor ion information). The precursor species were finally transferred and thermalized in segment Q5 by injecting a gas pulse in the Omnitrap platform before they were subjected to electron irradiation for 50 ms. Spectra were acquired over 10 min with the time-of-flight acquisition window recording over the 100-3000 *m/z* range. The Omnitrap platform is operated with the frequency of the rectangular waveform set at 800 kHz for optimal transmission and trapping efficiency of both precursor and fragment ions.

#### ExD fragmentation of native-like Fab precursor ions

Denatured Fabs were diluted in 150 mM ammonium acetate to a concentration of 5 μM, and loaded in aliquots of ~ 3 μL in in-house pulled emitters for nESI. Ions were accumulated for 400 ms in the TIMS cartridge, submitted to an in-source CID energy of 100 eV for desolvation, and then charge states 13+ and 14+ were mass isolated using the quadrupole mass filter with an isolation width of 10 Th (see **Supplementary Tables S1 and S2**). The Fab precursor species were finally transferred to segment Q5 and subjected to electron irradiation for 100 ms. Similarly, mass spectra were acquired over a period of 10 min in the 500-5000 *m/z* range with the time-of-flight acquisition window recording over the 100-3000 *m/z* range. The Omnitrap platform is operated with the frequency of the rectangular waveform set at 600 kHz for optimal transmission and trapping efficiency of both precursor and fragment ions.

It should be noted that, in all ExD experiments, the actual ion– electron reaction times are half of the values reported in this manuscript, as electrons are introduced into the Omnitrap platform only during one half of the rectangular waveform cycle.

### Data Analysis

Spectra were averaged over 10 min (for optimal signal quality) using Bruker Compass DataAnalysis version 6.1, and exported as *.xy data files for further processing in Bruker OmniScape 2025b version 1.10 with the “Confirmation” workflow (i.e., the workflow for fragment identification for proteins with a known sequence) to search for fragments against the known sequences of the individual Fab chains (Lc and Fd fragment, that were considered both with and without the mass of the counterpart chain as modification). Peak detection was performed with a simple moving average noise threshold, with a window size of 600 data points and a noise curve offset of 0.01. Fragment ion identification was then performed by looking for a-, b-, c-, x-, y- and z-type ions within a 5-ppm mass tolerance, a minimum intensity within mass tolerance of 70 %, a correlation threshold of 20 %, a score threshold of 4.0 a.u. and a stringency of peak acceptance set to “High”. All identifications were then manually inspected, and the resulting fragment ion lists were exported as *.csv files for further processing and visualization in R 4.4.0.

## ASSOCIATED CONTENT

### Supporting Information

- Description of Omniscape’s scoring system, ExD-MS/MS spectra and associated sequence maps for several charge states of 7D8-IgG1 and Trastuzumab Fabs, repartition of the fragments and illustrative zoom-ins on example fragments (PDF)
- Table of the timsOmni instrument parameters for analysis in denaturing and native conditions (XLSX)

## AUTHOR INFORMATION

### Author Contributions

The manuscript was written through contributions of all authors. All authors have given approval to the final version of the manuscript.

## ACKNOWLEDGMENT

A.J.R.H. acknowledges support from The Netherlands Organization for Scientific Research (NWO) through the Spinoza Award SPI.2017.028. We would like to thank Bruker Daltonics for support on this research. We thank Sjors P. A. van der Lans and Suzan H. M. Rooijakkers (Medical Microbiology, University Medical Center Utrecht, Utrecht, The Netherlands) for providing us with in-house produced recombinant IgdE.

## REFERENCES

(1) Bang, L. M.; Keating, G. M. Adalimumab. BioDrugs 2004, 18 (2), 121–139. 10.2165/00063030-200418020-00005.

(2) Mullard, A. FDA Approves 100th Monoclonal Antibody Product. Nat. Rev. Drug Discov. 2021, 20 (7), 491–495. 10.1038/d41573-021-00079-7.

(3) Rees, A. R. Understanding the Human Antibody Repertoire. mAbs 2020, 12 (1), 1729683. 10.1080/19420862.2020.1729683.

(4) Briney, B.; Inderbitzin, A.; Joyce, C.; Burton, D. R. Commonality despite Exceptional Diversity in the Baseline Human Antibody Repertoire. Nature 2019, 566 (7744), 393–397. 10.1038/s41586-019-0879-y.

(5) Krawczyk, K.; Kelm, S.; Kovaltsuk, A.; Galson, J. D.; Kelly, D.; Trück, J.; Regep, C.; Leem, J.; Wong, W. K.; Nowak, J.; Snowden, J.; Wright, M.; Starkie, L.; Scott-Tucker, A.; Shi, J.; Deane, C. M. Structurally Mapping Antibody Repertoires. Front. Immunol. 2018, 9. 10.3389/fimmu.2018.01698.

(6) Jiang, Y.; Rex, D. A. B.; Schuster, D.; Neely, B. A.; Rosano, G. L.; Volkmar, N.; Momenzadeh, A.; Peters-Clarke, T. M.; Egbert, S. B.; Kreimer, S.; Doud, E. H.; Crook, O. M.; Yadav, A. K.; Vanuopadath, M.; Hegeman, A. D.; Mayta, M. L.; Duboff, A. G.; Riley, N. M.; Moritz, R. L.; Meyer, J. G. Comprehensive Overview of Bottom-Up Proteomics Using Mass Spectrometry. ACS Meas. Sci. Au 2024, 4 (4), 338–417. 10.1021/acsmeasuresciau.3c00068.

(7) Yang, F.; Zhang, J.; Buettner, A.; Vosika, E.; Sadek, M.; Hao, Z.; Reusch, D.; Koenig, M.; Chan, W.; Bathke, A.; Pallat, H.; Lundin, V.; Kepert, J. F.; Bulau, P.; Deperalta, G.; Yu, C.; Beardsley, R.; Camilli, T.; Harris, R.; Stults, J. Mass Spectrometry-Based Multi-Attribute Method in Protein Therapeutics Product Quality Monitoring and Quality Control. mAbs 2023, 15 (1), 2197668. 10.1080/19420862.2023.2197668.

(8) Resemann, A.; Jabs, W.; Wiechmann, A.; Wagner, E.; Colas, O.; Evers, W.; Belau, E.; Vorwerg, L.; Evans, C.; Beck, A.; Suckau, D. Full Validation of Therapeutic Antibody Sequences by Middle-up Mass Measurements and Middle-down Protein Sequencing. mAbs 2016, 8 (2), 318–330. 10.1080/19420862.2015.1128607.

(9) Khristenko, N. A.; Nagornov, K. O.; Garcia, C.; Gasilova, N.; Gant, M.; Druart, K.; Kozhinov, A. N.; Menin, L.; Chamot-Rooke, J.; Tsybin, Y. O. Top-Down and Middle-Down Mass Spectrometry of Antibodies. Mol. Cell. Proteomics 2025, 24 (7), 100989. 10.1016/j.mcpro.2025.100989.

(10) Srzentić, K.; Fornelli, L.; Tsybin, Y. O.; Loo, J. A.; Seckler, H.; Agar, J. N.; Anderson, L. C.; Bai, D. L.; Beck, A.; Brodbelt, J. S.; van der Burgt, Y. E. M.; Chamot-Rooke, J.; Chatterjee, S.; Chen, Y.; Clarke, D. J.; Danis, P. O.; Diedrich, J. K.; D’Ippolito, R. A.; Dupré, M.; Gasilova, N.; Ge, Y.; Goo, Y. A.; Goodlett, D. R.; Greer, S.; Haselmann, K. F.; He, L.; Hendrickson, C. L.; Hinkle, J. D.; Holt, M. V.; Hughes, S.; Hunt, D. F.; Kelleher, N. L.; Kozhinov, A. N.; Lin, Z.; Malosse, C.; Marshall, A. G.; Menin, L.; Millikin, R. J.; Nagornov, K. O.; Nicolardi, S.; Paša-Tolić, L.; Pengelley, S.; Quebbemann, N. R.; Resemann, A.; Sandoval, W.; Sarin, R.; Schmitt, N. D.; Shabanowitz, J.; Shaw, J. B.; Shortreed, M. R.; Smith, L. M.; Sobott, F.; Suckau, D.; Toby, T.; Weisbrod, C. R.; Wildburger, N. C.; Yates, J. R.; Yoon, S. H.; Young, N. L.; Zhou, M. Interlaboratory Study for Characterizing Monoclonal Antibodies by Top-Down and Middle-Down Mass Spectrometry. J. Am. Soc. Mass Spectrom. 2020, 31 (9), 1783–1802. 10.1021/jasms.0c00036.

(11) Lavinder, J. J.; Horton, A. P.; Georgiou, G.; Ippolito, G. C. Next-Generation Sequencing and Protein Mass Spectrometry for the Comprehensive Analysis of Human Cellular and Serum Antibody Repertoires. Curr. Opin. Chem. Biol. 2015, 24, 112–120. 10.1016/j.cbpa.2014.11.007.

(12) Georgiou, G.; Ippolito, G. C.; Beausang, J.; Busse, C. E.; Wardemann, H.; Quake, S. R. The Promise and Challenge of High-Throughput Sequencing of the Antibody Repertoire. Nat. Biotechnol. 2014, 32 (2), 158–168. 10.1038/nbt.2782.

(13) de Graaf, S. C.; Hoek, M.; Tamara, S.; Heck, A. J. R. A Perspective toward Mass Spectrometry-Based de Novo Sequencing of Endogenous Antibodies. mAbs 2022, 14 (1), 2079449. 10.1080/19420862.2022.2079449.

(14) Rathore, A. S.; Sarin, D.; Bhattacharya, S.; Kumar, S. Multi-Attribute Monitoring Applications in Biopharmaceutical Analysis. J. Chromatogr. Open 2024, 6, 100166. 10.1016/j.jcoa.2024.100166.

(15) Carillo, S.; Criscuolo, A.; Füssl, F.; Cook, K.; Bones, J. Intact Multi-Attribute Method (iMAM): A Flexible Tool for the Analysis of Monoclonal Antibodies. Eur. J. Pharm. Biopharm. 2022, 177, 241–248. 10.1016/j.ejpb.2022.07.005.

(16) Bondt, A.; Dingess, K. A.; Hoek, M.; van Rijswijck, D. M. H.; Heck, A. J. R. A Direct MS-Based Approach to Profile Human Milk Secretory Immunoglobulin A (IgA1) Reveals Donor-Specific Clonal Repertoires With High Longitudinal Stability. Front. Immunol. 2021, 12, 789748. 10.3389/fimmu.2021.789748.

(17) Bondt, A.; Hoek, M.; Tamara, S.; Bastiaan, de G.; Peng, W.; Schulte, D.; van Rijswijck, D. M. H.; den Boer, M. A.; Greisch, J.-F.; Varkila, M. R. J.; Snijder, J.; Cremer, O. L.; Bonten, M. J. M.; Heck, A. J. R. Human Plasma IgG1 Repertoires Are Simple, Unique, and Dynamic. CELL Syst. 2021, 12 (12), 1131-+. 10.1016/j.cels.2021.08.008.

(18) Van Rijswijck, D. M. H.; Bondt, A.; Hoek, M.; Van Der Straten, K.; Caniels, T. G.; Poniman, M.; Eggink, D.; Reusken, C.; De Bree, G. J.; Sanders, R. W.; Van Gils, M. J.; Heck, A. J. R. Discriminating Cross-Reactivity in Polyclonal IgG1 Responses against SARS-CoV-2 Variants of Concern. Nat. Commun. 2022, 13 (1), 6103. 10.1038/s41467-022-33899-1.

(19) Fiala, J.; Schuster, D.; Ollivier, S.; Pengelley, S.; Lubeck, M.; Busch, F.; Jankevics, A.; Raether, O.; Greisch, J.-F.; Heck, A. J. R. Protein-Centric Analysis of Personalized Antibody Repertoires Using LC-MS-Based Fab-Profiling on a timsTOF. J. Am. Soc. Mass Spectrom. 2024, 35 (6), 1292–1300. 10.1021/jasms.4c00076.

(20) Brodbelt, J. S. Ion Activation Methods for Peptides and Proteins. Anal. Chem. 2016, 88 (1), 30–51. 10.1021/acs.analchem.5b04563.

(21) Brodbelt, J. S.; Morrison, L. J.; Santos, I. Ultraviolet Photodissociation Mass Spectrometry for Analysis of Biological Molecules. Chem. Rev. 2020, 120 (7), 3328–3380. 10.1021/acs.chemrev.9b00440.

(22) Greisch, J.-F.; Tamara, S.; A. Scheltema, R.; R. Maxwell, H. W.; D. Fagerlund, R.; C. Fineran, P.; Tetter, S.; Hilvert, D.; R. Heck, A. J. Expanding the Mass Range for UVPD-Based Native Top-down Mass Spectrometry. Chem. Sci. 2019, 10 (30), 7163–7171. 10.1039/C9SC01857C.

(23) Riley, N. M.; Coon, J. J. The Role of Electron Transfer Dissociation in Modern Proteomics. Anal. Chem. 2018, 90 (1), 40–64. 10.1021/acs.analchem.7b04810.

(24) Lermyte, F.; Valkenborg, D.; Loo, J. A.; Sobott, F. Radical Solutions: Principles and Application of Electron-Based Dissociation in Mass Spectrometry-Based Analysis of Protein Structure. Mass Spectrom. Rev. 2018, 37 (6), 750–771. 10.1002/mas.21560.

(25) Lantz, C.; Schrader, R.; Meeuwsen, J.; Shaw, J.; Goldberg, N. T.; Tichy, S.; Beckman, J.; Russell, D. H. Digital Quadrupole Isolation and Electron Capture Dissociation on an Extended Mass Range Q-TOF Provides Sequence and Structure Information on Proteins and Protein Complexes. J. Am. Soc. Mass Spectrom. 2023, 34 (8), 1753–1760. 10.1021/jasms.3c00184.

(26) Baba, T.; Ryumin, P.; Duchoslav, E.; Chen, K.; Chelur, A.; Loyd, B.; Chernushevich, I. Dissociation of Biomolecules by an Intense Low-Energy Electron Beam in a High Sensitivity Time-of-Flight Mass Spectrometer. J. Am. Soc. Mass Spectrom. 2021, 32 (8), 1964–1975. 10.1021/jasms.0c00425.

(27) Chen, X.; Wang, Z.; Wong, Y.-L. E.; Wu, R.; Zhang, F.; Chan, T.-W. D. Electron-Ion Reaction-Based Dissociation: A Powerful Ion Activation Method for the Elucidation of Natural Product Structures. Mass Spectrom. Rev. 2018, 37 (6), 793–810. 10.1002/mas.21563.

(28) Zubarev, R. A. Electron-Capture Dissociation Tandem Mass Spectrometry. Curr. Opin. Biotechnol. 2004, 15 (1), 12–16. 10.1016/j.copbio.2003.12.002.

(29) Mao, Y.; Valeja, S. G.; Rouse, J. C.; Hendrickson, C. L.; Marshall, A. G. Top-Down Structural Analysis of an Intact Monoclonal Antibody by Electron Capture Dissociation-Fourier Transform Ion Cyclotron Resonance-Mass Spectrometry. Anal. Chem. 2013, 85 (9), 4239–4246. 10.1021/ac303525n.

(30) Fornelli, L.; Damoc, E.; Thomas, P. M.; Kelleher, N. L.; Aizikov, K.; Denisov, E.; Makarov, A.; Tsybin, Y. O. Analysis of Intact Monoclonal Antibody IgG1 by Electron Transfer Dissociation Orbitrap FTMS *. Mol. Cell. Proteomics 2012, 11 (12), 1758–1767. 10.1074/mcp.M112.019620.

(31) Shaw, J. B.; Liu, W.; Vasil′ev, Y. V.; Bracken, C. C.; Malhan, N.; Guthals, A.; Beckman, J. S.; Voinov, V. G. Direct Determination of Antibody Chain Pairing by Top-down and Middle-down Mass Spectrometry Using Electron Capture Dissociation and Ultraviolet Photodissociation. Anal. Chem. 2020, 92 (1), 766–773. 10.1021/acs.analchem.9b03129.

(32) Greisch, J.-F.; den Boer, M. A.; Beurskens, F.; Schuurman, J.; Tamara, S.; Bondt, A.; Heck, A. J. R. Generating Informative Sequence Tags from Antigen-Binding Regions of Heavily Glycosylated IgA1 Antibodies by Native Top-Down Electron Capture Dissociation. J. Am. Soc. Mass Spectrom. 2021, 32 (6), 1326–1335. 10.1021/jasms.0c00461.

(33) Greisch, J.-F.; den Boer, M. A.; Lai, S.-H.; Gallagher, K.; Bondt, A.; Commandeur, J.; Heck, A. J. R. Extending Native Top-Down Electron Capture Dissociation to MDa Immunoglobulin Complexes Provides Useful Sequence Tags Covering Their Critical Variable Complementarity-Determining Regions. Anal. Chem. 2021, 93 (48), 16068–16075. 10.1021/acs.analchem.1c03740.

(34) Fornelli, L.; Srzentić, K.; Huguet, R.; Mullen, C.; Sharma, S.; Zabrouskov, V.; Fellers, R. T.; Durbin, K. R.; Compton, P. D.; Kelleher, N. L. Accurate Sequence Analysis of a Monoclonal Antibody by Top-Down and Middle-Down Orbitrap Mass Spectrometry Applying Multiple Ion Activation Techniques. Anal. Chem. 2018, 90 (14), 8421–8429. 10.1021/acs.analchem.8b00984.

(35) Cotham, V. C.; Brodbelt, J. S. Characterization of Therapeutic Monoclonal Antibodies at the Subunit-Level Using Middle-Down 193 Nm Ultraviolet Photodissociation. Anal. Chem. 2016, 88 (7), 4004–4013. 10.1021/acs.analchem.6b00302.

(36) Cotham, V. C.; Horton, A. P.; Lee, J.; Georgiou, G.; Brodbelt, J. S. Middle-Down 193-Nm Ultraviolet Photodissociation for Unambiguous Antibody Identification and Its Implications for Immunoproteomic Analysis. Anal. Chem. 2017, 89 (12), 6498–6504. 10.1021/acs.analchem.7b00564.

(37) Butalewicz, J. P.; Escobar, E. E.; Wootton, C. A.; Theisen, A.; Park, M. A.; Seeley, E. H.; Brodbelt, J. S. Conformational Characterization of Peptides and Proteins by 193 Nm Ultraviolet Photodissociation in the Collision Cell of a Trapped Ion Mobility Spectrometry-Time-of-Flight Mass Spectrometer. Anal. Chem. 2024, 96 (41), 16154–16161. 10.1021/acs.analchem.4c02686.

(38) Fung, Y. M. E.; Adams, C. M.; Zubarev, R. A. Electron Ionization Dissociation of Singly and Multiply Charged Peptides. J. Am. Chem. Soc. 2009, 131 (29), 9977–9985. 10.1021/ja8087407.

(39) Papanastasiou, D.; Kounadis, D.; Lekkas, A.; Orfanopoulos, I.; Mpozatzidis, A.; Smyrnakis, A.; Panagiotopoulos, E.; Kosmopoulou, M.; Reinhardt-Szyba, M.; Fort, K.; Makarov, A.; Zubarev, R. A. The Omnitrap Platform: A Versatile Segmented Linear Ion Trap for Multidimensional Multiple-Stage Tandem Mass Spectrometry. J. Am. Soc. Mass Spectrom. 2022, 33 (10), 1990–2007. 10.1021/jasms.2c00214.

(40) Zhai, Z.; Mavridou, D.; Damian, M.; Mutti, F. G.; Schoenmakers, P. J.; Gargano, A. F. G. Characterization of Complex Proteoform Mixtures by Online Nanoflow Ion-Exchange Chromatography-Native Mass Spectrometry. Anal. Chem. 2024, 96 (22), 8880–8885. 10.1021/acs.analchem.4c01760.

