## supplemental data for "A prototype timsOmni platform enables confident annotation of key hypervariable regions of IgG immunoglobulins using low- and high-energy electron-based fragmentation"

**On the calculations of scores using OmniScape**

For a given cleavage site, a score with a value between 0 and 1, representing a probability, is associated to every matched fragment ion. For a given cleavage site, the cumulative distribution function (CDF) of all the scores of all the matched fragments gives you the probability of detecting a fragment with a score value less than or equal to x. For a given score x, the product of all probabilities (roughly, under certain assumptions) gives you the probability of having a set of fragments with score values less than x. For a given amino acid, the total score is the product of the scores associated with all the cleavages immediately (before the next amino acid) to the left and to the right of the amino acid considered. If the value is close to 1, it means that the fragment combination corresponds to a high-confidence amino acid assignment. If the value is close to 0, it means that the fragment combination provides a lower-confidence assignment. The blue coloring scheme used in main **Figure 6** corresponds to a linear gradient where 0 is no color and 1 is totally blue.

**Supplementary Table S1.** Detailed timsOmni experimental instrument parameters used for the analysis of the Fabs ionized under denaturing and native MS conditions.

*See attached xlsx file*

**Supplementary Table S2.** Precursor ions *m/z* and Q0 isolation width used.

| | Charge state | 7D8-IgG1 | | Trastuzumab | |
|---|---|---|---|---|---|
| | | Precursor *m/z* | Isolation width (*m/z*) | Precursor *m/z* | Isolation width (*m/z*) |
| Denatured charge states | **31+** | 1546.2 | 5 | 1533.3 | 5 |
| | **34+** | 1409.9 | 5 | 1398.0 | 5 |
| | **37+** | 1295.6 | 5 | 1284.8 | 5 |
| Native-like charge states | **13+** | 3685.7 | 10 | 3654.8 | 10 |
| | **14+** | 3422.5 | 10 | 3393.8 | 10 |

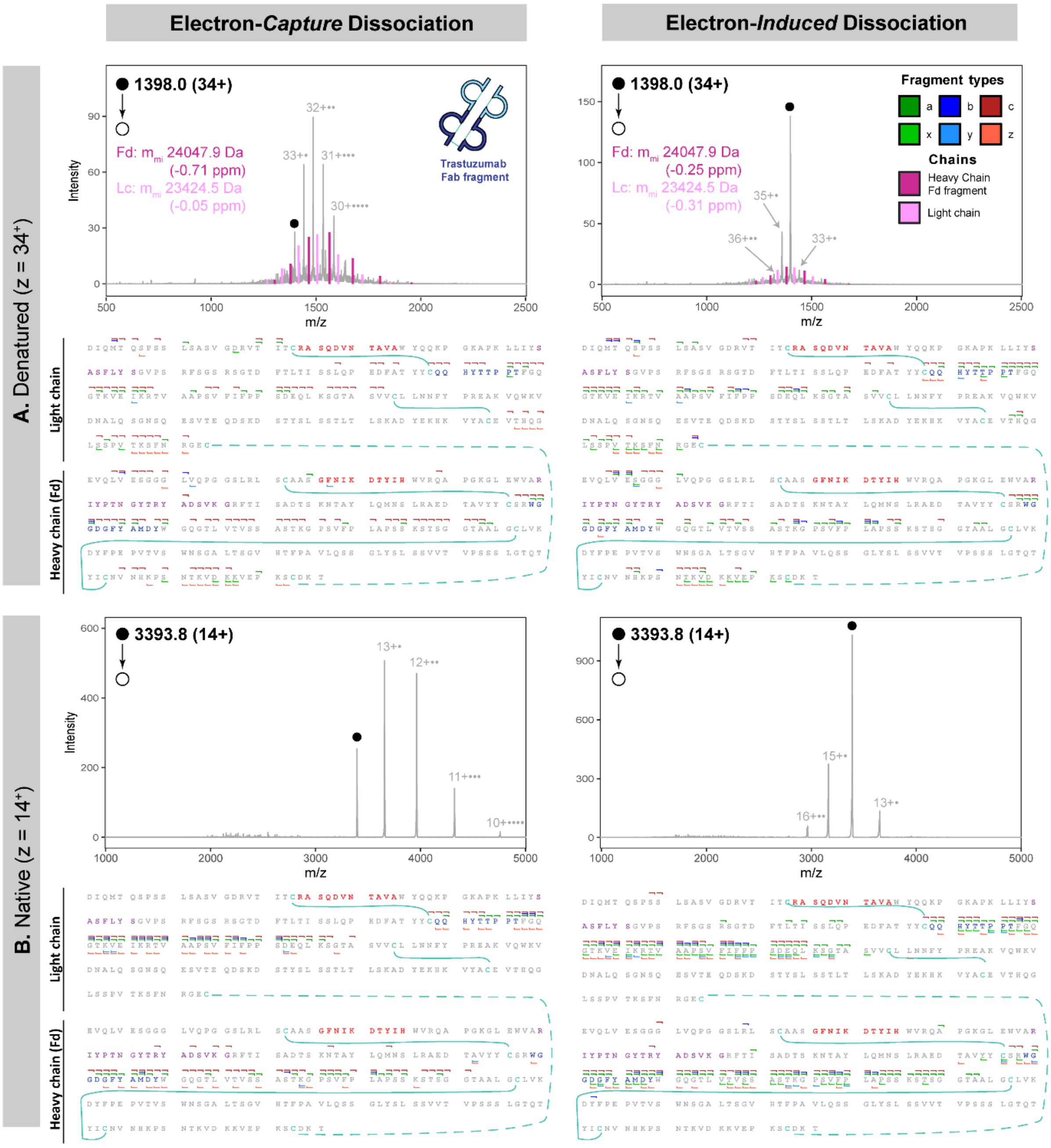

**Figure S1. ExD provides information on Lc/Fd pairing and sequence coverage of the CDR3 region, as in Figure 4, shown here for Trastuzumab.** MS2 ExD spectra of the Trastuzumab Fab and confidently annotated fragment ions mapped onto its sequence, with the CDR1, CDR2 and CDR3 highlighted in red, purple and blue respectively for **(A)** higher charge state Fab ions produced under denaturing conditions (z=34+), and **(B)** lower charge state Fab ions produced under native conditions (z=14+) both subjected to low-energy ECD (left) and to high-energy EID (right).

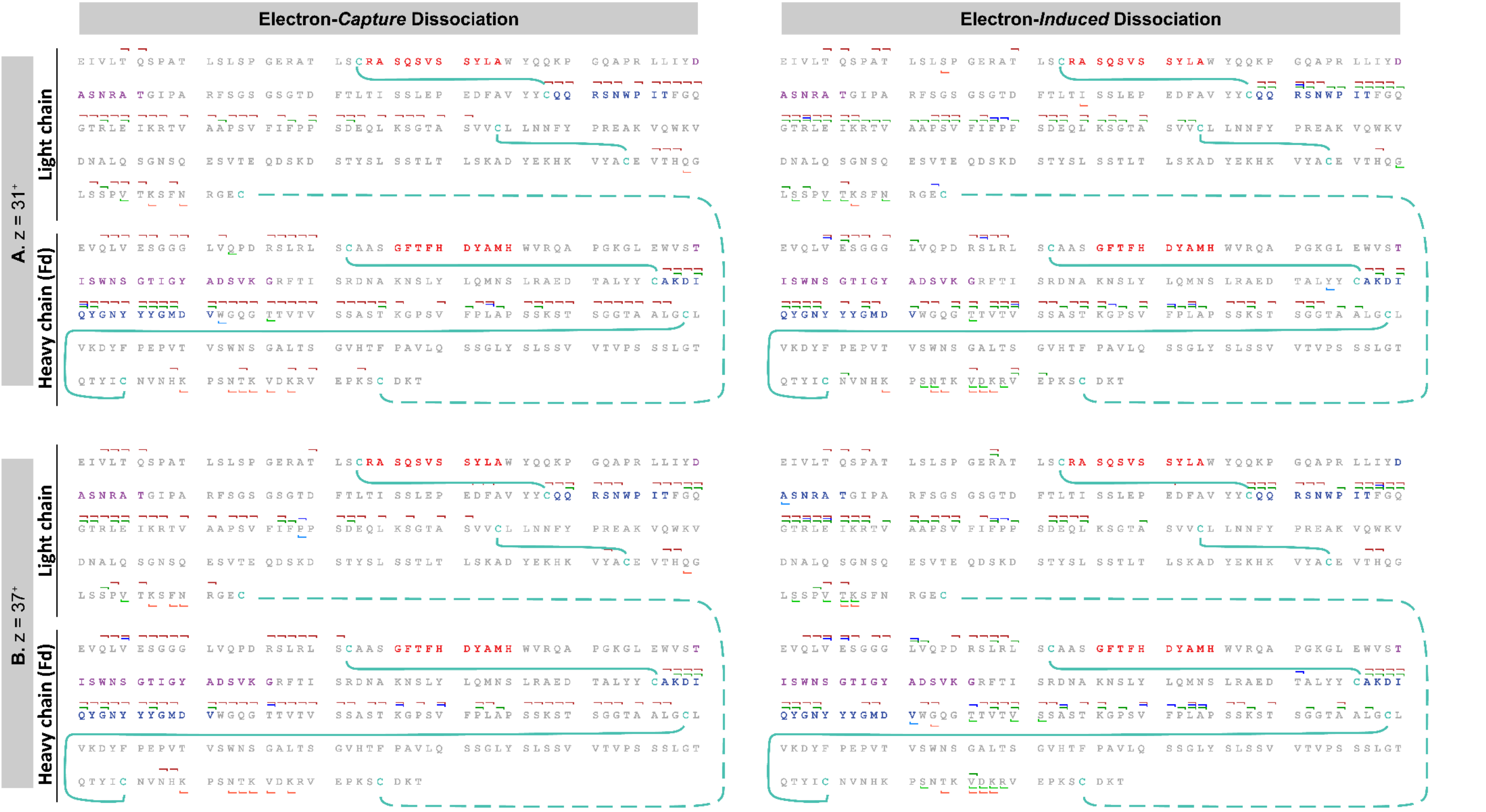

**Figure S2. The charge states of precursor ions have low impact on the fragmentation of Fabs produced under denaturing conditions.** Confidently annotated fragment ions from MS2 ExD of 7D8-IgG1 Fab are mapped onto its sequence, with the CDR1, CDR2 and CDR3 highlighted in red, purple and blue respectively for **(A)** z=31$^{+}$ and **(B)** z=37$^{+}$ higher charge state Fab ions subjected to low-energy ECD (left), and high-energy EID (right).

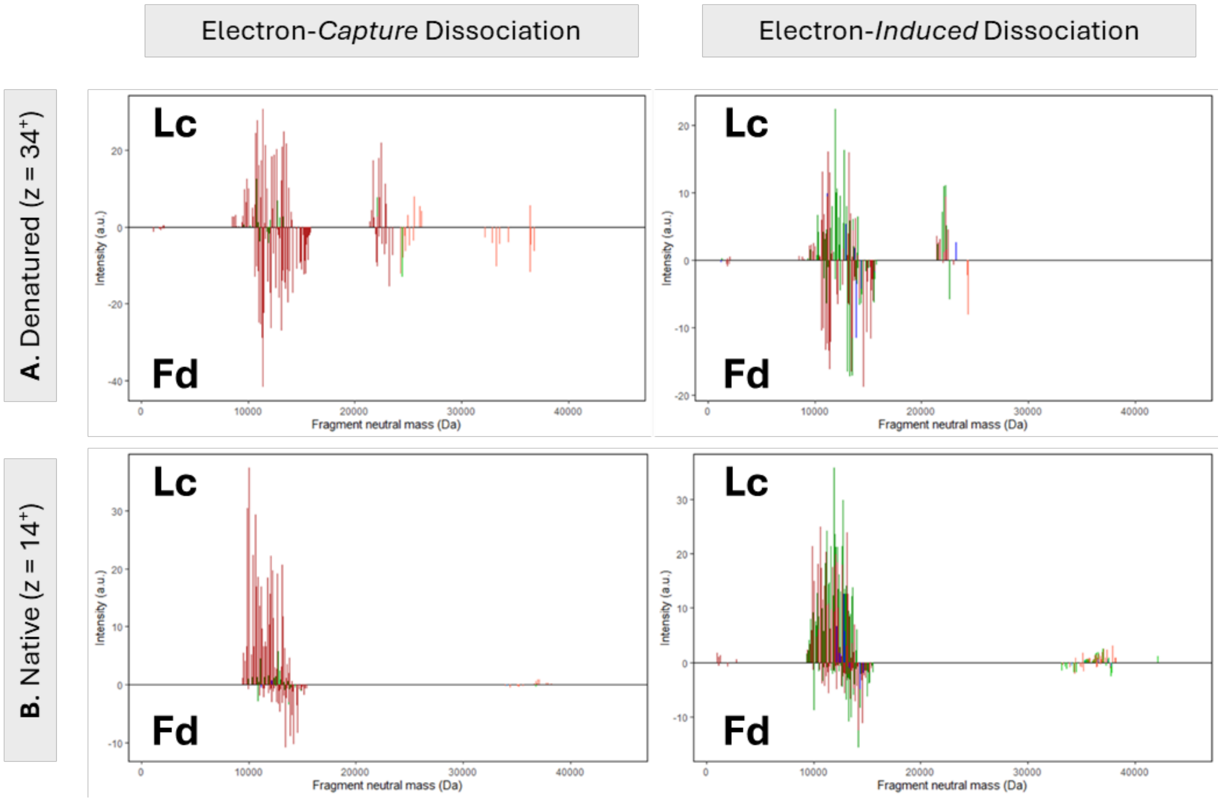

**Figure S3. When Fabs are produced under denaturing conditions (higher charge state precursors), the fragments from both chains have similar intensities, but when produced under native conditions (lower charge state precursors) the fragments from the Fd region of the Hc are less intense.** Neutral mass plots show the distributions of the fragments from both chains of **(A)** z=34+ 7D8-IgG1 Fab and **(B)** z=14+ 7D8-IgG1 Fab in ECD (left) and EID (right).

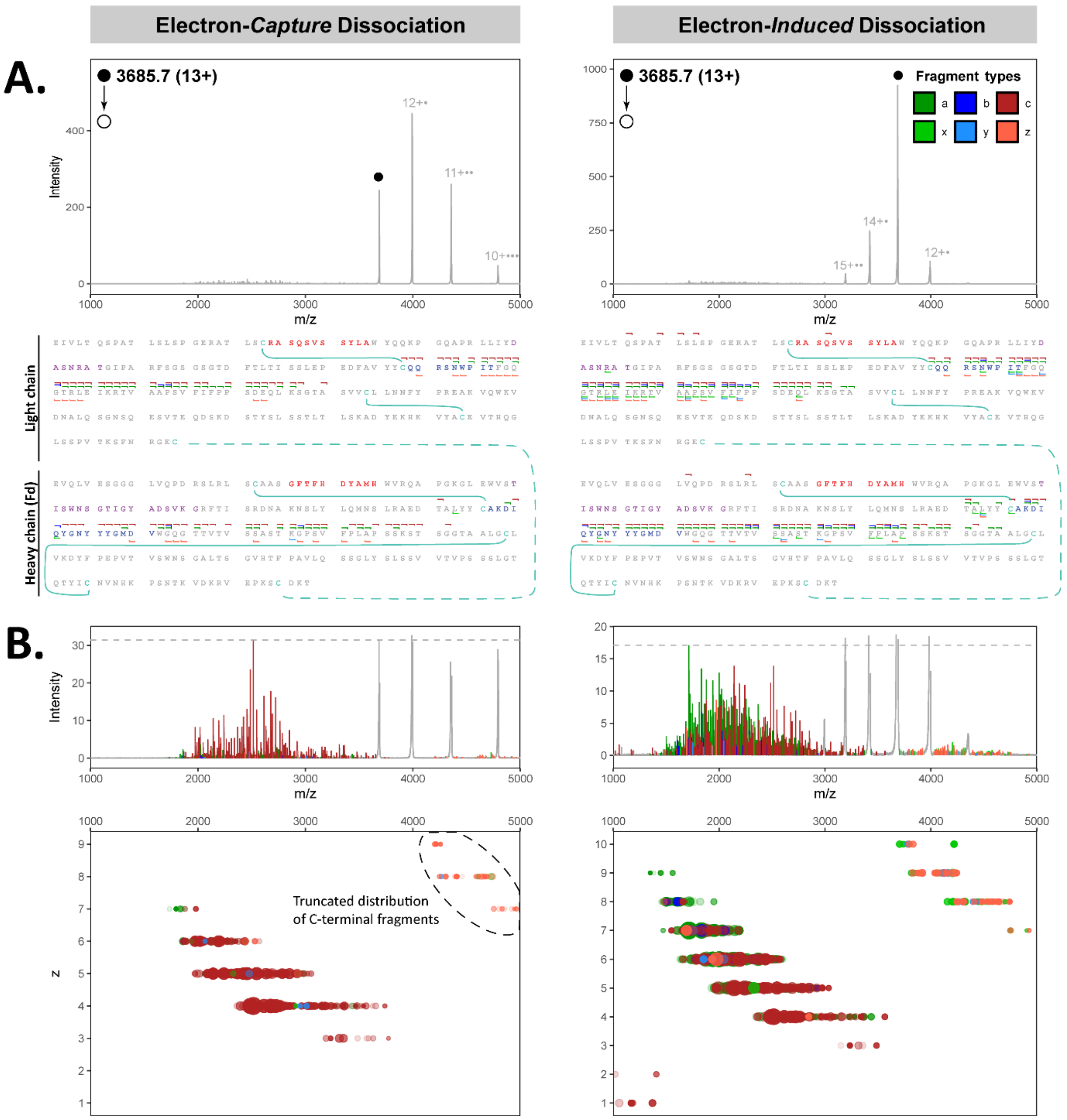

**Figure S4. Impact of the precursor charge state in MS2 ExD of Fab molecules produced under native conditions. (A) MS2** ExD spectra (ECD on the left, EID on the right) of the z=13+ charge state of 7D8-IgG1 Fab and confidently annotated fragment ions mapped onto its sequence, with the CDR1, CDR2 and CDR3 highlighted in red, purple and blue respectively. The coverage by C-terminal fragments is reduced compared to the 14+ charge state of 7D8-IgG1 (main text **Figure 4B**), especially in the case of ECD. **(B)** A zoomed y-axis view of the annotated fragment ions and corresponding charge versus *m/z* plot show that the lower coverage by C-terminal ions for the 13+ Fab arises from these fragments carrying a lower number of charges, causing their distribution to shift beyond the *m/z* range of the analysis.

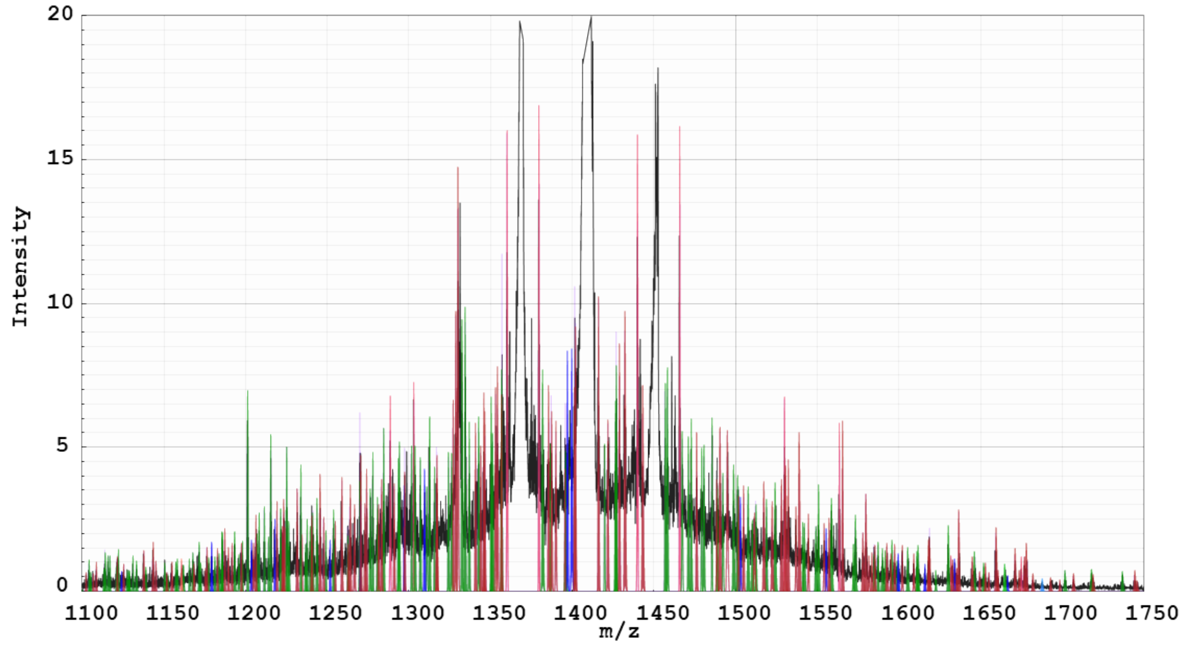

**Figure S5. An extreme overlap of fragment ions and charge-increased precursor distributions occurs in EID of Fab higher charge states produced under denaturing conditions.** A zoom-in on the *m/z* 1000-2000 range of the MS2 EID spectrum of the 34+ 7D8-IgG1 Fab shows the overlap between fragment ions and charge-increased precursor species.

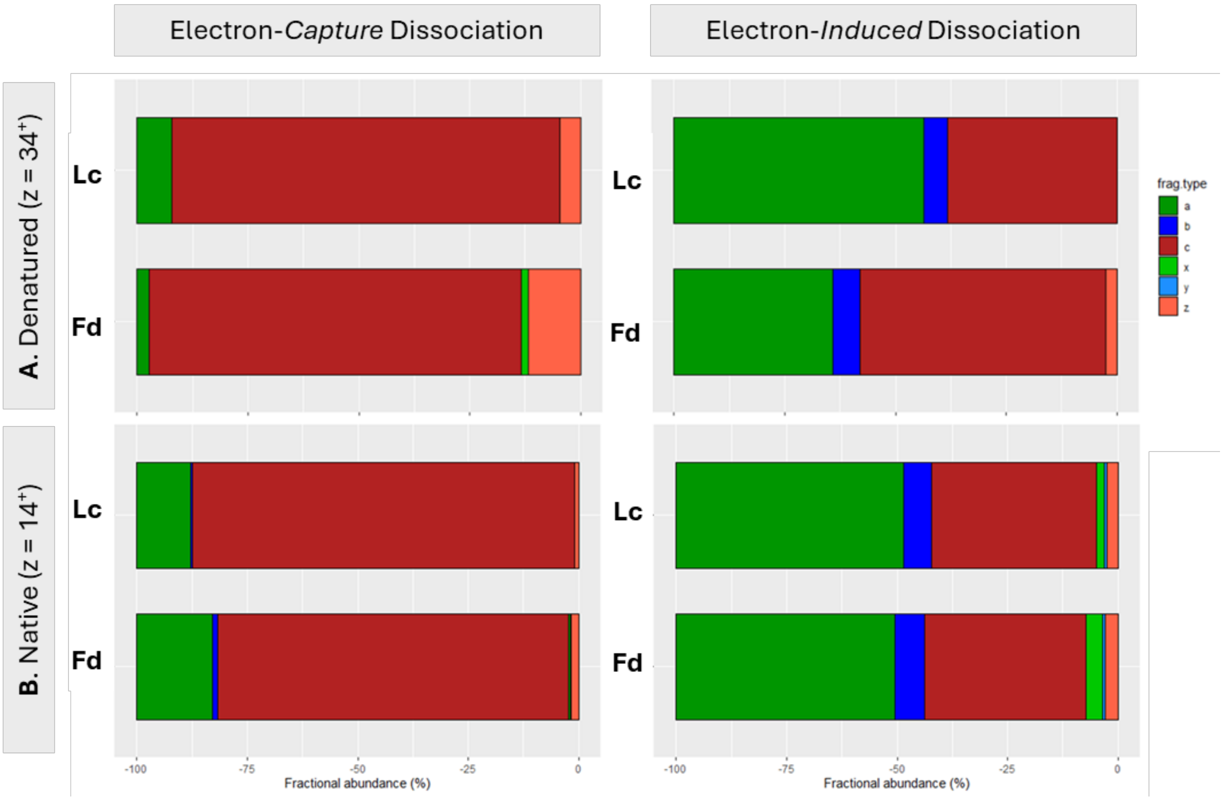

**Figure S6. Intensity-based fractional abundance plots of the different types of fragments** generated from (A) z=34+ charge state 7D8-IgG1 Fab produced under denaturing conditions, and (B) z=14+ charge state 7D8-IgG1 Fab produced under native-like conditions during ECD (left) and EID (right). Significantly more a-type fragment ions, along with a low number of *b*-type ions, are observed in EID compared to ECD. Under denaturing conditions, C-terminal fragments are largely indistinguishable due to extensive spectral overlap, whereas native conditions facilitate the identification of C-terminal x-, y- and z-type ions.

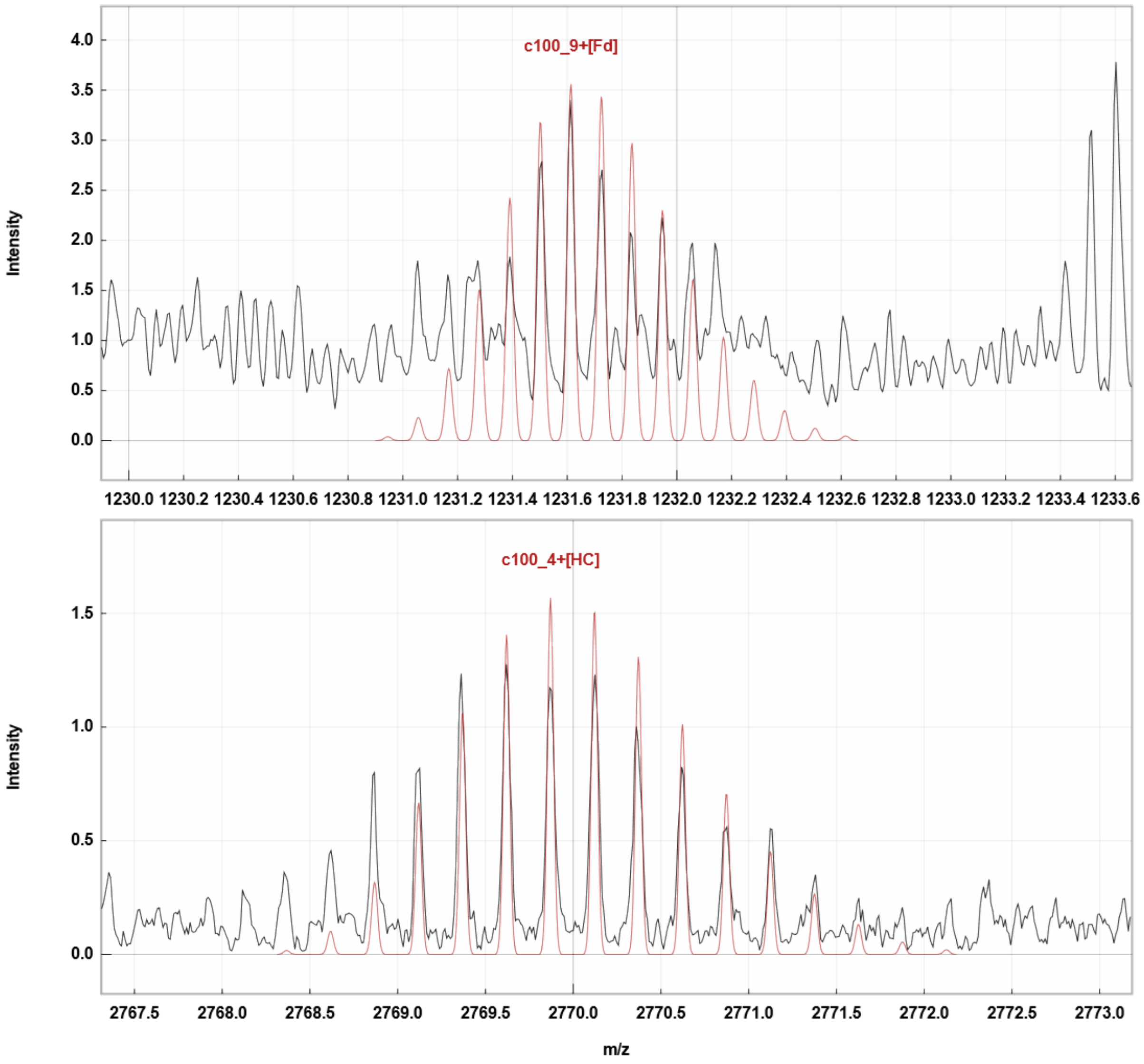

**Figure S7. The lower charge states of fragments obtained from Fabs produced under native conditions reduce spectral congestion, facilitating their identification compared to those measured under denaturing conditions.** This is exemplified by the c100 fragment of the heavy chain observed following EID of the z=34+ (top) and z=14+ (bottom) 7D8-IgG1 Fab precursors.

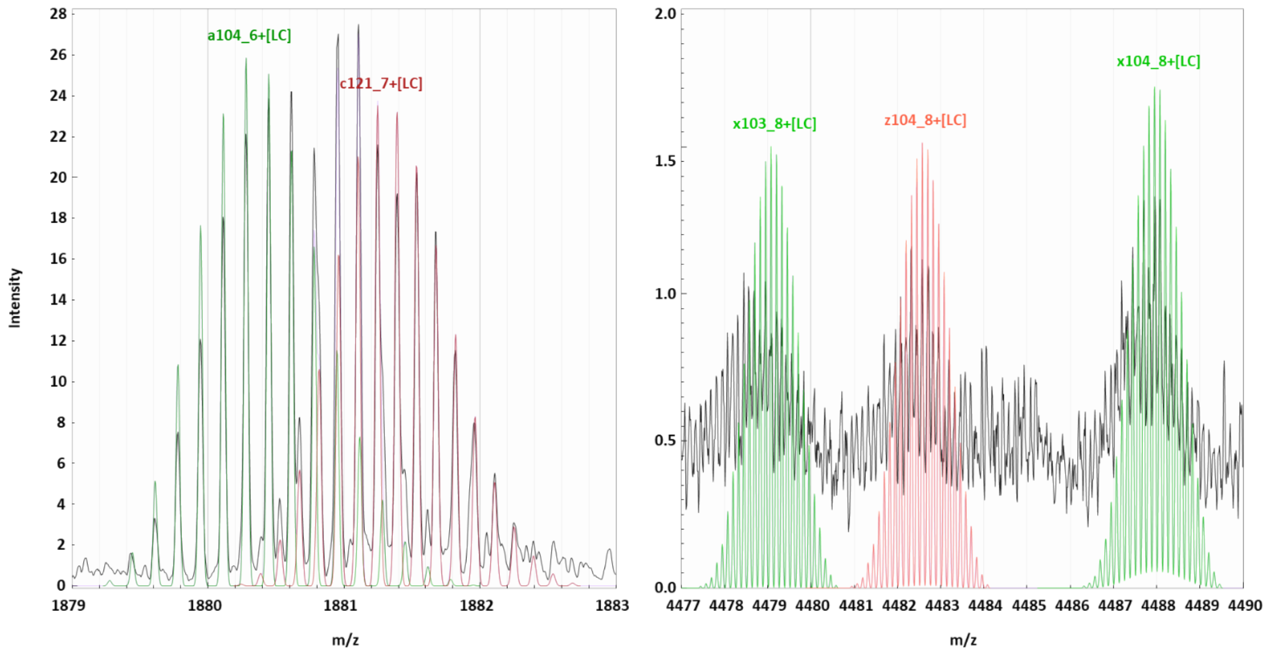

**Figure S8. Isotopic profile comparison of the N- and C-terminal fragments generated from Fabs analysed under native conditions**. N-terminal a- and c- fragment ions (left) containing part of a single chain, and C-terminal x- and z- ions (right) retaining the counterpart chain are illustrated on the MS2 EID spectrum of z=14+ 7D8-IgG1 Fab. The C-terminal ions are only partially resolved and exhibit broadened isotopic distributions, resulting in lower S/N ratios.

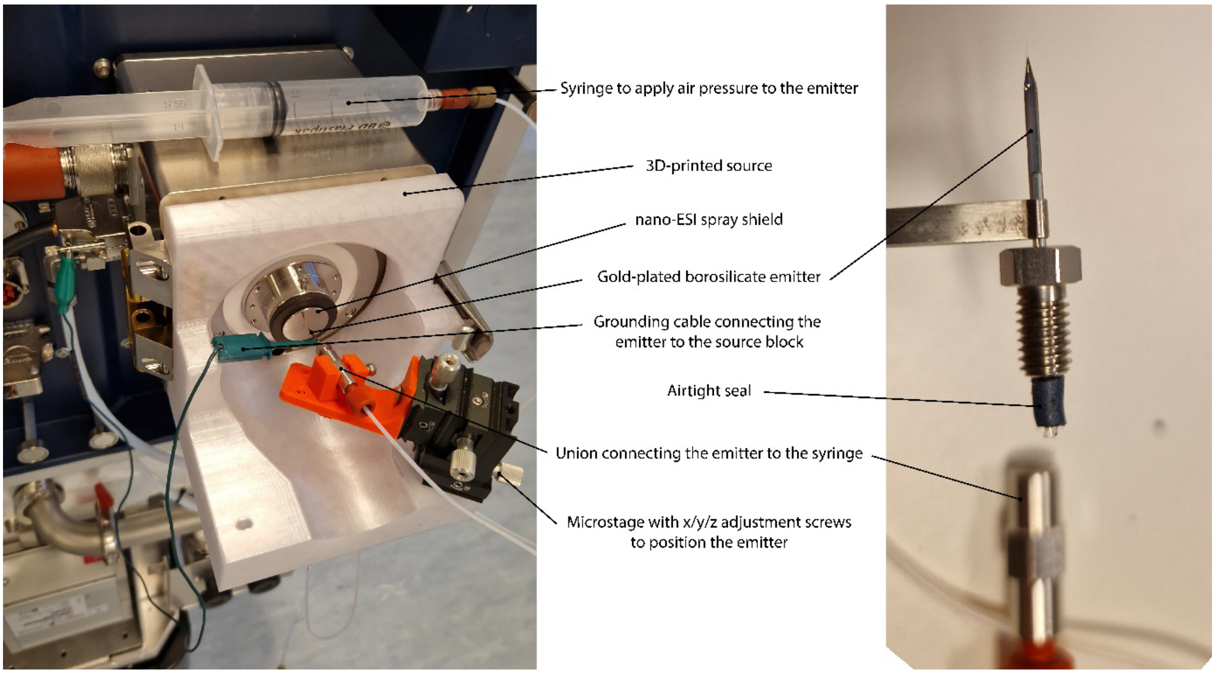

**Figure S9. In-house-designed nanoESI ionization source for the timsOmni**, used with in-house produced gold-plated borosilicate emitters.
